# Convergent Evolution During Local Adaptation to Patchy Landscapes

**DOI:** 10.1101/006940

**Authors:** Peter L. Ralph, Graham Coop

**Affiliations:** Computational Biology and Bioinformatics, University of Southern California, Los Angeles, CA; Center for Population Biology & Department of Evolution and Ecology, University of California – Davis, Davis, CA, 95616

## Abstract

Species often encounter, and adapt to, many patches of locally similar environmental conditions across their range. Such adaptation can occur through convergent evolution if different alleles arise and spread in different patches, or through the spread of shared alleles by migration acting to synchronize adaptation across the species. The tension between the two reflects the degree of constraint imposed on evolution by the underlying genetic architecture versus how effectively selection and geographic isolation act to inhibit the geographic spread of locally adapted alleles. This paper studies a model of the balance between these two routes to adaptation in continuous environments with patchy selection pressures. We address the following questions: How long does it take for a novel, locally adapted allele to appear in a patch of habitat where it is favored through mutation? Or, through migration from another, already adapted patch? Which is more likely to occur, as a function of distance between the patches? How can we tell which has occurred, i.e., what population genetic signal is left by the spread of migrant alleles? To answer these questions we examine the family structure underlying migration–selection equilibrium surrounding an already adapted patch, in particular treating those rare families that reach new patches as spatial branching processes. This provides a way to understand the role of geographic separation between patches in promoting convergent adaptation and the genomic signals it leaves behind. We illustrate these ideas using the convergent evolution of cryptic coloration in the rock pocket mouse, *Chaetodipus intermedius*, as an empirical example.

**Author Summary:** Often, a large species range will include patches where the species differs because it has adapted to locally differing conditions. For instance, rock pocket mice are often found with a coat color that matches the rocks they live in, these color differences are controlled genetically, and mice that don’t match the local rock color are more likely to be eaten by predators. Sometimes, similar genetic changes have occurred independently in different patches, suggesting that there were few accessible ways to evolve the locally adaptive form. However, the genetic basis could also be shared if migrants carry the locally beneficial genotypes between nearby patches, despite being at a disadvantage between the patches. We use a mathematical model of random migration to determine how quickly adaptation is expected to occur through new mutation and through migration from other patches, and study in more detail what we would expect successful migrations between patches to look like. The results are useful for determining whether similar adaptations in different locations are likely to have the same genetic basis or not, and more generally in understanding how species adapt to patchy, heterogeneous landscapes.

## Introduction

The convergent evolution of similar phenotypes in response to similar selection pressures is a testament to the power that selection has to sculpt phenotypic variation. In some cases this convergence extends to the molecular level, with disparate taxa converging to the same phenotype through parallel genetic changes in the same pathway, genes, or even by precisely the same genetic changes [Stern, 2013, Zhen et al., 2012, Martin and Orgogozo, 2013]. Convergent adaptation also occurs within species, if different individuals adapt to the same environment through different genetic changes. There are a growing number of examples of this in a range of well studied organisms and phenotypes [Arendt and Reznick, 2008]. Such evolution of convergent phenotypes is favored by higher mutational input, i.e., higher total mutational rate and/or population size [Pennings and Hermisson, 2006]. The geographic distribution of populations can also affect the probability of parallel mutation within a species: a widespread species is more likely to adapt by multiple, parallel mutations if dispersal is geographically limited, since subpopulations will adapt via new mutations before the adaptive allele arrives via migration [Ralph and Coop, 2010]. Standing variation for a trait can also increase the probability of convergence, as this increases the probability that the selective sweep will be *soft* (beginning from a base of multiple copies), which leads to genetic patterns similar to convergent adaptation [Orr and Betancourt, 2001, Hermisson and Pennings, 2005] whether or not the copies derive from the same mutation,

Intuitively, convergence is also more likely when geographically separated populations adapt to ecologically similar conditions. The probability that convergent adaptations arise independently before adaptations spread between the populations by migration will be larger if these adaptive alleles are maladapted in intervening environments, since such adverse conditions can strongly restrict the spread of locally adapted alleles [Slatkin, 1973].

One elegant set of such examples is provided by the assortment of plant species that have repeatedly adapted to patches of soil with high concentrations of heavy metals (e.g., serpentine outcrops and mine tailings) [Kruckeberg, 1951, Macnair, 1991, Schat et al., 1996, Turner et al., 2010]; the alleles conferring heavy metal tolerance are thought to be locally restricted because they incur a cost off of these patches. Similarly, across the American Southwest, a variety of species of animals have developed locally adaptive cryptic coloration to particular substrates, e.g., dark rock outcrops or white sand dunes [Benson, 1933]. One of the best-known examples is the rock pocket mouse (*Chaetodipus intermedius*), which on a number of black lava flows has much darker pelage than on intervening areas of lighter rock [Dice and Blossom, 1937]. Strong predator-mediated selection appears to favour such crypsis [Kaufman, 1974], and, perhaps as a result of this strong selection against migrants, at least two distinct genetic changes are responsible for the dark pigmentation found on different outcrops [Hoekstra and Nachman, 2003]. Similar situations have been demonstrated in other small rodent systems [Dice, 1940, Steiner et al., 2009, Kingsley et al., 2009] and in lizards [Rosenblum et al., 2010].

In this paper, we study this situation, namely, when a set of alleles provide an adaptive benefit in geographically localized patches, separated by inhabited regions where the alleles are deleterious. The main questions are: Under what conditions is it likely that each patch adapts in parallel, i.e., convergently through new mutation, and when is it likely that migration carries these alleles between patches? How easy will it be to detect adaptive alleles that are shared by migration, i.e., over what genomic scale will a population genetic signal be visible?

We work in a model of continuous geography, using a variety of known results and new methods. In the section *Establishment by Mutation* we develop a simple approximation, equation (2), for the rate at which patches become established by new mutations. The most important conceptual work of the paper occurs in the section *Establishment by Migration*, where we develop an understanding of the process by which alleles move from an existing patch to colonize a novel patch, culminating in equation (11) for the rate at which this happens. We combine these two results in the section *Probability of Parallel Adaptation* to discuss the probability of parallel adaptation, equation (12). To understand the genomic signal left by adaptations shared by migration, in the section *Haplotypes Shared Between Patches*, we study the time it will take for an allele to transit between patches, (equation (18)), and thus the length of haplotype that hitchhikes with it (equation (19)). Finally, in the section *Applications* we apply this work to understand the geographic scale of convergent evolution in *Chaetodipus intermedius*.

## Results

### Model of a patchy landscape

Consider a population spread across a landscape to which it is generally well adapted, but within which are patches of habitat to which individuals are (initially) poorly adapted. (When we refer to “patches” it is to these pieces of poor habitat.) Suppose it takes only a single mutational change to create an allele (*B*) that adapts an individual to the poor habitat type. The required change could be as specific as a single base change, or as general as a knockout of a gene or one of a set of genes. This change occurs at a (low) rate of *μ* per chromosome per generation, and has fitness advantage *s*_*p*_ relative to the unmutated type (*b*) in these “poor” habitat patches. Outside of these patches the new allele has a fitness disadvantage of *s*_*m*_ in the intervening areas, with *s*_*p*_ and *s*_*m*_ both positive. (Here we define “fitness” to be the intrinsic growth rate of the type when rare.) We assume that the disadvantage *s*_*m*_ is sufficiently large that, on the relevant timescale, the allele is very unlikely to fix locally in these intervening regions. (The case where *s*_*m*_ = 0 requires a different approach, which we do not treat here.)

We are interested in the establishment of mutations in the “poor” patches by either migration or mutation, which depends on whether the allele can escape initial loss by drift when rare. Therefore, we need not specify the fitness of the homozygote; only that the dynamics of the allele when rare are determined by the fitness of the heterozygote. More general dominance will only make small corrections to the dynamics until initial fixation, with the exception of the recessive case, which we omit. In other words, we follow the literature in treating the diploid model as essentially haploid.

We also assume population density *ρ* is constant across the range (even in the “poor” patches). The variance in offspring number is *ξ*^2^, and that the mean squared distance between parent and child is *σ*^2^ (i.e., *σ* is the dispersal distance). We will deal with migration by immediately appealing to the central limit theorem, treating the total distance traveled across *t* dispersal events as Gaussian with variance *tσ*^2^, and do not discuss the (interesting) cases where rare, long-distance dispersal events are more important (see e.g., Levin et al. [2003], Ralph and Coop [2010], Hallatschek and Fisher [2014] for discussion).

#### Past work on spatial models

We will make use of several previous results from the literature on migration in models of spatially varying selection. Haldane [1948], Fisher [1950], and Slatkin [1973] first analyzed deterministic models of of this sort. Nagylaki [1975] and Conley [1975] showed that in one dimension, if the physical width of the patch is less than 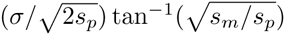, then there is no stable equilibrium with both *b* and *B* present, so that migrational swamping prevents *B* from establishing (see also Lenormand [2002] for a review). Barton [1987], using general theory of Pollak [1966], showed that patches must be at least this critical size so that a new mutation has positive probability of establishment (in the large population density limit). The same paper also showed that this probability of establishment of a new mutation decays exponentially with distance from the patch, with the scale given by 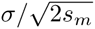. Mutations appearing within the patch have probability of establishment strictly less than the probability for a panmictic population, but approaching this as the size of the patch increases. This latter panmictic probability of establishment we denote *p*_*e*_, and often approximate *p*_*e*_ by 2*s*_*p*_*/* ξ ^2^ [Haldane, 1927, Fisher, 1930]. This result holds quite generally for a geographically spread population that experiences a uniform selection pressure [Maruyama, 1970, Cherry and Wakeley, 2003]. More general work on the mathematically equivalent problem of density-dependent population growth has obtained much more general criteria under which a habitat configuration will support a stable polymorphic equilibrium [Cantrell and Cosner, 1989], and determined the habitat configuration that maximizes the probability of establishment [Lou and Yanagida, 2006].

### Establishment of a locally adaptive allele due to mutational influx

We first compute the time scale on which a new *B* mutations will appear and fix in a single, isolated poor habitat patch in which no *B* allele has yet become established. As we are interested in patches where local adaptation can occur, we will assume that the patch is larger than the cutoff for local establishment mentioned above.

Let *p*(*x*) be the probability that a new mutant *B* allele that arises at location *x* relative to the center of the patch fixes within the poor habitat patch. Under various assumptions, precise expressions can be found for *p*(*x*) [Barton, 1987], but results will be more interpretable if we proceed with a simple approximation. The total influx per generation of mutations that successfully establish is the product of population density *ρ*, mutation rate *μ*, and the integral of *p*(*x*) over the entire species range:

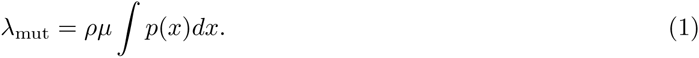

This depends in a complicated way on the patch geometry and selection coefficients, but still scales linearly with the mutational influx density *ρμ*. If we consider a patch of area *A*, whose width is large relative to 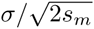, then a reasonable approximation is to ignore migration, so that *p*(*x*) = *p*_*e*_ *≈* 2*s*_*p*_/ξ^2^ within the patch, and *p*(*x*) = 0 otherwise. This approximates the integrand *p*(*x*) in (1) by a step function, which will be a good approximation if the patch is large relative the scale over which *p*(*x*) goes from 0 to *p*_*e*_, or if *p*(*x*) is close to *p*_*e*_ at some point and is symmetric about the edge of the patch. We examine this more generally via exact calculation of *p*(*x*) in the section *Probability of Establishment*.

The rate at which mutations arise and colonize a patch of area *A* is then

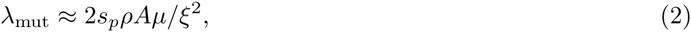

i.e., the product of mutational influx in the patch (*ρAμ*) and the probability of establishment (*p*_*e*_ *≈* 2*s*_*p*_*/ξ*^2^). If this rate is low, then the time (in generations) until a mutation arises that will become locally established within the patch is exponentially distributed with mean 1*/λ*_mut_. Assuming that once a mutation becomes established it quickly reaches its equilibrium frequency across the patch, the time scale on which new patches become colonized by the *B* allele from new mutation is therefore approximately 1*/λ*_mut_.

### Establishment of a locally adaptive allele due to migrational influx

Now suppose that there are two patches separated by distance *R* (i.e., the shortest distance between the two is *R*). If the *B* allele has arisen and become established in the first patch, but has not yet appeared in the second, we would like to know the time scale on which copies of *B* move between patches by migration (supposing that no *B* allele arises independently by mutation in the meantime). To determine this, we study the fine-scale genealogy of alleles that transit between patches, obtaining along the way other useful information about the genealogy of *B* alleles. Doing this we arrive at equation (11) for the rate at which an allele established in one patch spreads to a neighboring one by migration.

Migration–selection balance ensures that there will be some *B* alleles present in the regions where they are disadvantageous, but only rarely, far away from the patch where *B* is established. Denote the expected frequency of allele *B* at displacement *x* relative to the patch by *q*(*x*), and assume that the shape of the patch is at least roughly circular. Following Haldane [1948] or Slatkin [1973], one can write down a differential equation to which *q*(*x*) is the solution, and show that *q*(*x*) decays exponentially for large *|x|*, with a square-root correction in two dimensions:

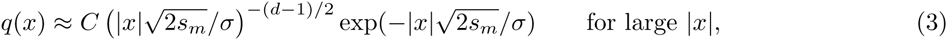

where *d* is the dimension (*d* = 1 or 2), and *C* is a constant depending on the geographic shape of the populations and the selection coefficients. In applications we fit *C* to the data; to get concrete numbers from theory we take *C* = 1 if necessary.

As J. Hermisson pointed out in comments on an earlier draft, this functional form has a simple intuition: local migration leads to the exponential decay, since if each migrant at distance *x* produces an average of *c* descendants that make it to distance *x* + 1, then in one dimension, the number of migrants must decay as exp(*-cx*); and the square-root term in two dimensions enters because the available area grows with *x*. The “renewal” aspect of this same argument suggests that for equation (3) to hold, *|x|* must be larger than a few multiples of *σ*.

These asymptotics fit simulations quite well, as shown in Figure 1. To be clear about the assumptions implicit here, below we provide a derivation of (3) in section *Equilibrium Frequency*, as well as justification for the asymptotics below in section *Asymptotics*. In one dimension, the equation can be solved to give *q*(*x*) in terms of Jacobi elliptic functions; see the Supporting Information.

**Figure 1.**
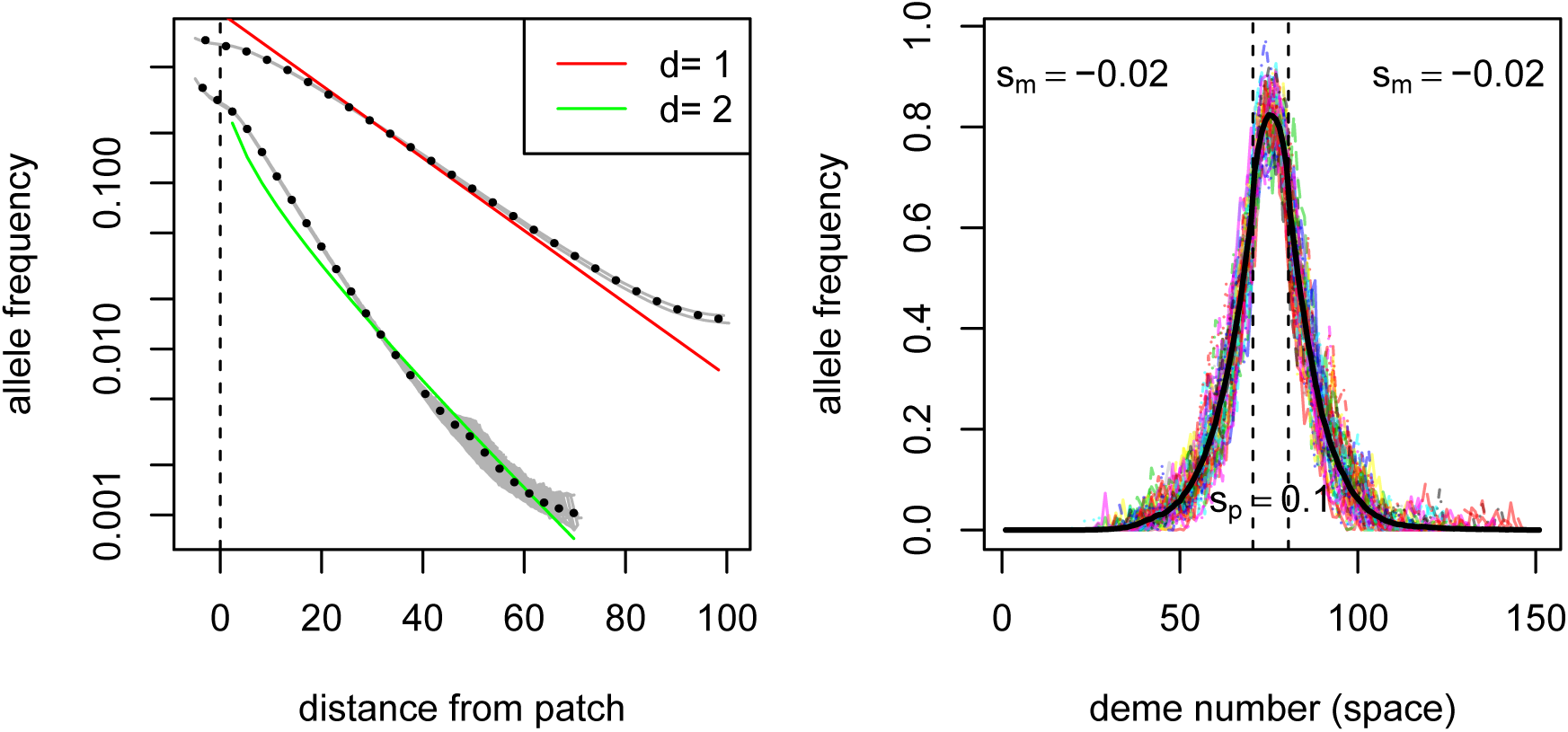
**(A)** Mean occupation frequencies of individual demes (grey dots) and averaged over demes at similar distances (black circles) from simulations, on a one-dimensional grid of 201 demes, and a two-dimensional grid of 101 *×* 101 demes. Superimposed in color is the decay predicted by (3), with *C* chosen arbitrarily to give a reasonable fit: 0.85 and 0.11 respectively. (The disagreement at long distances is due to boundary effects near the edge of the range.) **(B)** Fluctuations: colored lines are snapshots of allele frequencies at evenly spaced times from a one-dimensional grid of 151 demes; the solid line shows the cumulative mean occupation frequency across all generations. For both plots, the simulation was run for 1000 generations; the focal allele is beneficial in the central 10 demes, with *s*_*p*_ = 0.1, and deleterious elsewhere, with *s*_*m*_ = −0.02; demes had 1000 individuals in each; dispersal was to nearby demes with mean dispersal distances of *σ* = 0.95 and 0.74 times the deme spacing, respectively. More details are given below in *Simulation methods*.

This expected frequency *q*(*x*)is the *time-averaged* occupation frequency, i.e., the total number of *B* alleles found across *T* generations near location *x*, per unit area, divided by *T*. If, for instance, *q*(*x*) = .01 and the population density is *ρ* = 10 individuals per unit area, then in a survey tract of unit area around *x* we expect to find one individual every 10 generations – or, perhaps more likely, 10 individuals every 100 generations. This points to an important fact: close to the original patch, the “equilibrium frequency” *q*(*x*) describes well the density of the *B* allele at most times, but far away from the patch, the equilibrium is composed of rare excursions of families of *B* alleles, interspersed by long periods of absence. An example of one of these rare excursions is shown in Figure 2. The relevant time scale on which *B* alleles migrate between patches is given by the rate of appearance of such families.

**Figure 2.**
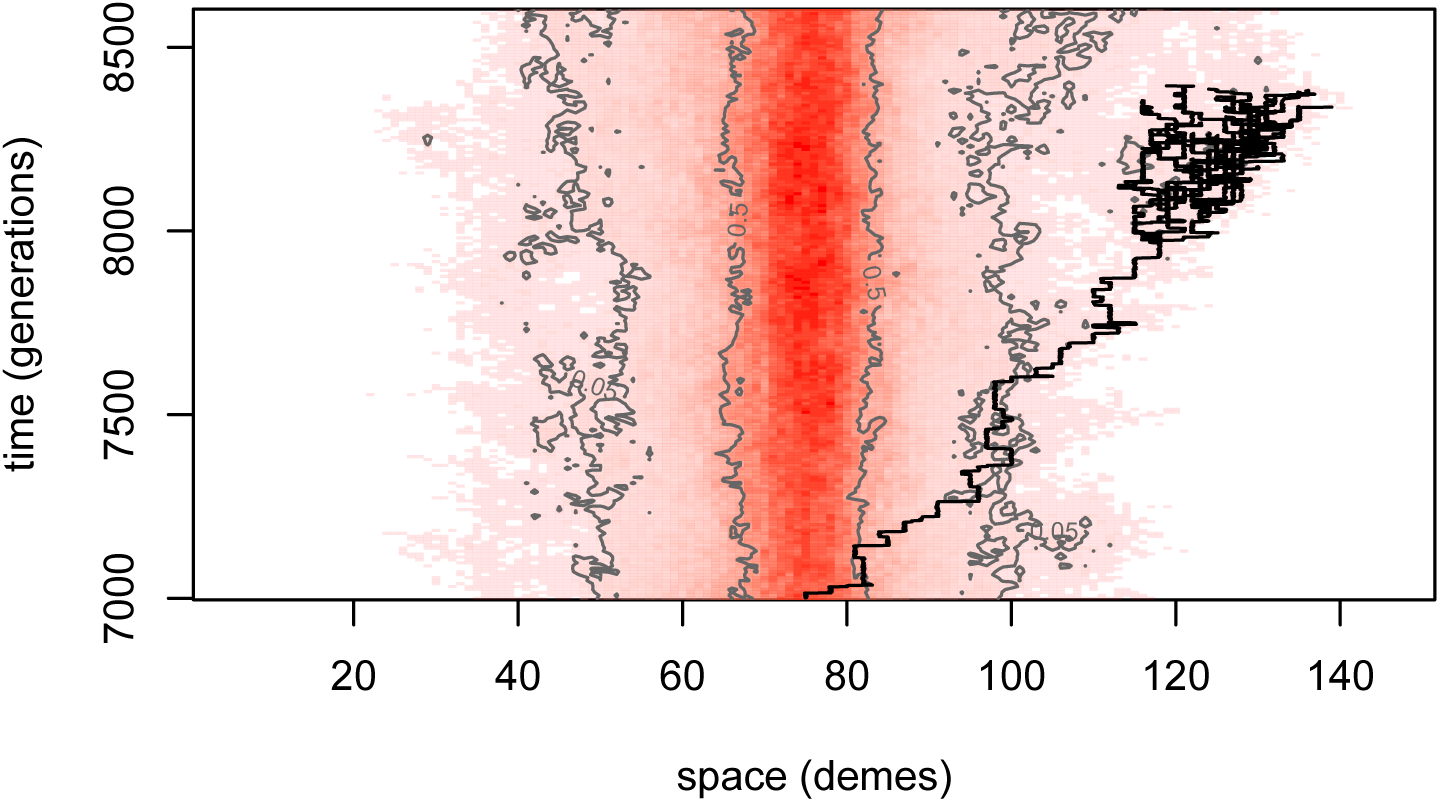
A segment of the temporal dynamics of the same simulation as Figure 1B. Space is again shown across the horizontal axis, with time on the vertical axis; demes are colored darker red the greater their allele frequency at the corresponding time. The 50% and 5% frequency contour lines are in grey, and the genealogy of 50 individuals in a rare long-distance excursion is traced back with black lines.

This suggests decomposing the genealogy of *B* alleles into families in the following way. First, consider the genealogy of all *B* alleles that were alive at any time outside of the original patch. This is a collection of trees, each rooted at an allele living outside the patch whose parent was born inside the patch. Next, fix some intermediate distance *r*_0_ from the established patch, and erase from the genealogy every allele that has no ancestors living further away than *r*_0_ to the patch. This fragments the original set of trees into a collection of smaller trees that relate to each other all *B* alleles living outside of *r*_0_ at any time, and some living inside of *r*_0_; we refer to these trees as “families”. If *r*_0_ is large enough that there are not too many families and interactions between family members are weak, then these families can be approximated by a collection of independent spatial branching processes whose members are ignored if they enter the original patch, illustrated in figure 3. (This can be made formal in the limit of large population density, also taking *r*_0_ large enough that the chance of reaching the original patch is small.) The opportunity for adaptation depends on how often these families of *B* alleles encounter the new patch. Suppose that the area occupied by the new patch is *S*. We can learn about the rate of arrival of families at *S* from the time-averaged number of *B* alleles expected in *S* were it not a patch (i.e., if *B* was still maladaptive in *S*), which from equation (3) is

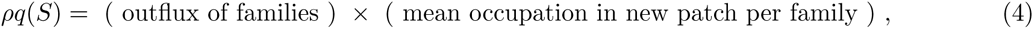

where 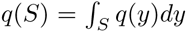. The quantity we wish to compute, the effective rate at which *families of* migrant *B* alleles establish in the new patch, is

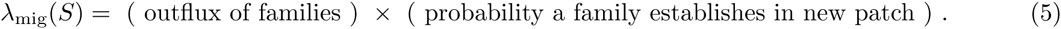

The quantities on the right-hand side depend implicitly on *r*_0_, but their product does not.

### The genealogy of migrant families

To understand the process of adaptation by migration, we therefore need to look more closely at the rare families of *B* alleles far from the original patch. It is reasonable to model each such family as a branching process, i.e., assuming that each individual migrates, then reproduces, independently of all other family members. Migration takes the individual in a random direction for a random distance whose mean square is *σ*^2^, and if the individual arrives to a patch where the *B* allele is favored, we drop the individual from the family. Since the *B* allele is uniformly deleterious elsewhere, we can decouple reproduction and migration by first forming a tree in which each individual has a random number of offspring, then assigning spatial locations to each individual in the tree, and finally pruning any branches that wander into a patch. This pruning removes both branches leading back into the already established patch (which are irrelevant) and branches leading into the new patch (we will need to count these). Since we are concerned with the rare migrant families that transit quickly between patches, we ignore back migrants to the already established patch.

Since each individual has on average 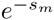 offspring, the expected family size after time *t* is 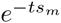. Therefore, if we define *k*_*e*_(*t*) to be the chance that the family has gone extinct by time *t*, and let *K*_*t*_ be the (random) family size conditioned on nonextinction, then by Bayes’ rule, 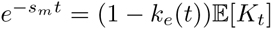. It turns out that if the distribution of offspring numbers is not too heavy-tailed (see Jagers [1975] for details), then *K*_*t*_ has a limiting distribution: 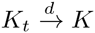, and

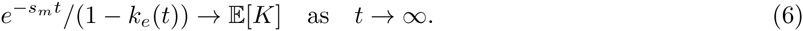

In other words, the chance of survival to time *t* is asymptotically a constant multiplied by 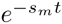, and conditional on survival, the family size is given by a fixed distribution.

This suggests that families can be well-described by motion of a central “trunk”, which dies at a random time whose distribution is given by *k*_*e*_(*t*), surrounded by “branches”, as depicted in Figure 3. We can understand this with a decomposition described by Geiger [1999]. When we condition on survival until *t*, we condition on survival of at least one trunk lineage (the long red lineage in Figure 3). Given this lineage, the other reproduction events are undistinguished – the genealogy of a family that survives until *t* can be decomposed into a trunk that lasts until *t* and that sprouts independent, unconditioned branches that may or may not survive until *t*. Concretely, if we write *Z*_*t*_ for the number of offspring in the branching process alive at time *t*, then

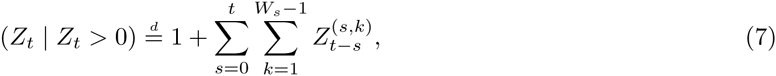

where each *Z*^(*s*,*k*)^ is an independent copy of *Z* that arose from the main branch in generation *s*, and each *W*_*s*_ is independent with distribution that tends to ℙ{*W*_*s*_ = *k*} ℙ{*W = k*}*/*𝔼[*W*] as *s → ∞*, where *W* is the unconditioned offspring number distribution. In particular, 𝔼[*W*_*s*_ − 1] 𝔼[*W* (*W -* 1)]*/*𝔼[*W*]; if *W* is Poisson then this is equal to 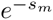. Since each subfamily *Z*^(*s*,*k*)^ is a branching process with mean offspring number less than one, most of these die out quickly, and so *Z*_*t*_ is composed of a cluster of recently split subfamilies. If we take the expectation of (7), since 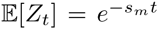, we get the limiting mean family size conditioned on nonextinction:

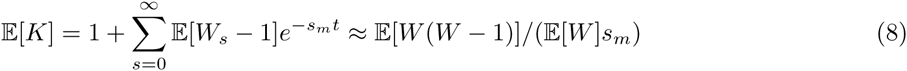

**Figure 3.**
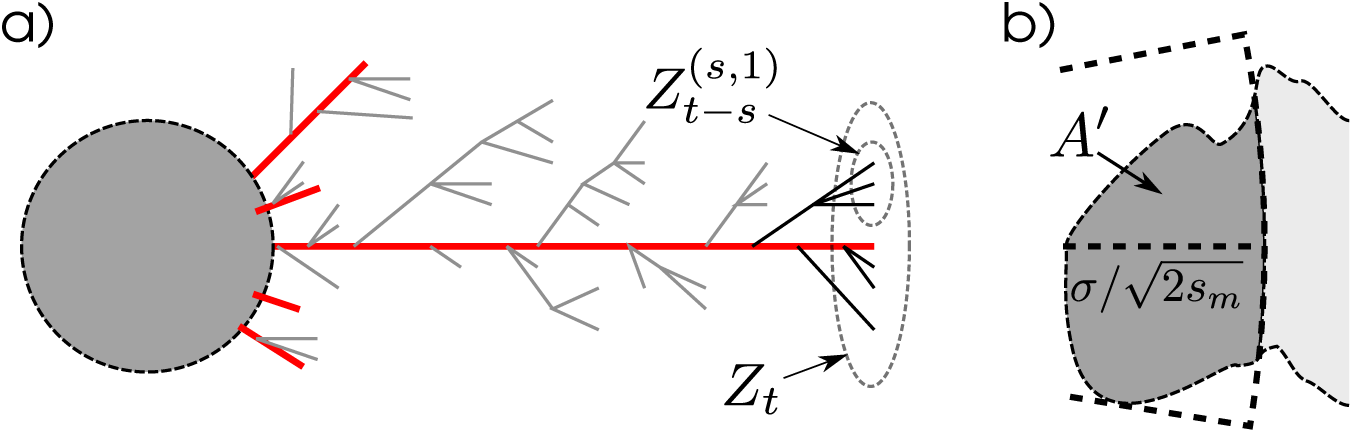
**(a)** Cartoon of the decomposition described in the text. Migrants families that get further than *r*_0_ from the already-established patch (grey circle) are each composed of a “trunk” lineage (in red), whose spatial motion we follow, embellished by transient branches. For the depicted long-lived family, these are further distinguished by those still alive at time *t* (in black) and those that did not survive until *t* (in grey). Figure 2 has a depiction of such a family, from a simulation. Note that the spatial motions of lineages are independent random walks (and hence should be much wigglier than in this cartoon, which confounds space and time). **(b)** Diagram of the area *A*′ used in equation (11): *R* is the shortest distance from the old patch to the new, and *A*′ is the area of the portion of the new patch that is no more than distance 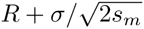 from the old patch.

Subfamilies typically live for 1*/s*_*m*_ generations, thus spreading across an area of width 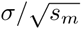. The oldest subfamily in a surviving migrant family is of order log(1*/s*_*m*_)*/s*_*m*_ generations, so the entire family is spread slightly wider.

### Heuristics

The parallel form suggested by (4) and (5) suggests that we can find *λ*_mig_(*S*) by multiplying the mean density *q*(*S*) by the ratio of the second terms in the two equations, i.e.

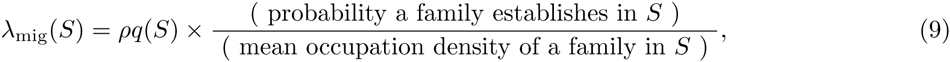

where again the “occupation density” is computed *without* the selective advantage. To make this precise we need to understand what we mean by “outflux of families”; we do this below in section *Hitting and Occupation*, and provide a heuristic argument here.

Both numerator and denominator in (9) count only families that have arrived at the new patch, so we can consider a family conditioned on having successfully transited between the patches. To obtain simply interpretable expressions, we assume that once any member of the family arrives at the patch, all other members are close by, and so have roughly equal opportunity to establish in the new patch.

Suppose the family arrives with *K* individuals. The probability that at least one individual leaves a lineage that establishes in the patch is 1 − (1 − *p*_*e*_)^*K*^. On the other hand, in the absence of the patch of new habitat, each of the *K* newly arrived individuals would leave behind a lineage with mean total size 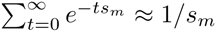, roughly half of which would stay in the new patch. Therefore, the expected contribution of the family to the occupation density is *K/*2*s*_*m*_. (For more careful consideration of the geometry of the patch and the distinction between occupation density and number of individuals, see section *Hitting and Occupation*, below.)

This suggests that

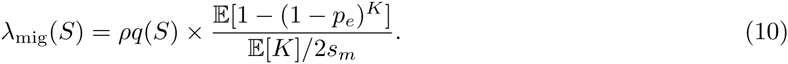

There are now two extremes to consider. If ℙ{*K >* 1*/p*_*e*_} is small, then each family has a small chance of establishment, and the numerator is approximately *p*_*e*_𝔼[*K*]. On the other hand, if *p*_*e*_ is not too small, then the numerator would be close to 1, as any family that arrives will likely establish.

The remaining term is *q*(*S*), the integral of equation (3) over the patch. Since the patches are at distance *R*, if the new patch has width smaller than 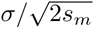, then *q*(*S*) is approximately *q*(*R*) multiplied by the area of the new patch. If *S* is larger, then we should use the area of the closest strip of width 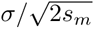 to the original patch; see figure 3, and section *The Integral q*(*S*) for more details. In either case, denote this area *A*′ ; for numerical examples we compute *q*(*S*) directly.

Combining these, and absorbing the term E[*W* (*W -* 1)]*/*E[*W*] from equation (8) into the constant *C*,

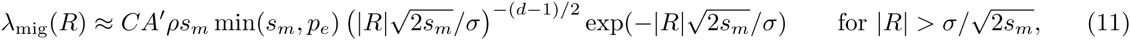

where *A*′ is the area of the new patch no further than 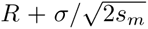 from the old patch. When we use this below, we set *p*_*e*_ = 2*s*_*p*_*/ξ*^2^. Note that the approximation has the undesirable property that it is non-monotonic in *s*_*m*_, when we expect the rate of adaptation via migration to be a decreasing function of *s*_*m*_. However, it only increases for small *s*_*m*_, and is a decreasing function of *s*_*m*_ for *s*_*m*_ *> |R|*^2^*/*2*σ*^2^, which is the parameter region that we are restricted to by other assumptions.

### A review, and a test, of the assumptions

Now that we have expressions for the mean rates of adaptation by new mutation, (2), and by migration from an already colonized patch, (11), it seems helpful to step back and review the assumptions underlying the asymptotic results we have used, or will use below. Our results should hold exactly in the limit of large, circular patches sufficiently far apart, large population density, and small selection coefficients of equal magnitude.

To be more precise, the mutation rate results most obviously apply if *s*_*m*_ ≈ *s*_*p*_, if the patch diameter is large relative to 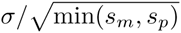, and the patch is not too far from circular. If *s*_*m*_ ≪ *s*_*p*_ then a strip of width 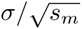 should be added to the outside of the patch in computing *A* for equation (2), and if *s*_*m*_ *≫ s*_*p*_, a strip of width 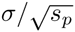 should be subtracted from the patch. As for the migration rate, we assume that each patch is large enough to support a stable population of *B* alleles 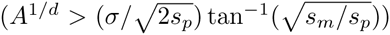. The geometry and size of the patch will also affect the approximation of equation (48). Next, in using equation(3), we assume that the inter-patch distance is large relative to the characteristic length 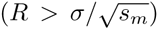, and that local drift is not so strong that the equilibrium frequency is at least approximately attained (using Wright’s effective neighborhood size, *s*_*m*_ *≫* 1*/*(*ρσ*^*d*^)). The last requirement is necessary because if the *B* allele fixes in large neighborhoods where it is deleterious, we cannot approximate its dynamics via a branching process. We also neglect the time for migration-selection equilibrium to be reached. As discussed above, we also assume that migration to a new patch takes place over a number of generations; if there are sufficient rare, long-distance migration events that would move between patches in a single hop, this would require a different analysis.

To test the robustness of our results to the various approximations we used, we used individual-based simulations on a one-dimensional lattice of 501 demes, with *ρ* haploid individuals per deme, dispersal to nearby demes with *σ* = 0.95, and run for 25,000 generations. More details are given below in section *Simulation methods*, and the number of simulations used and parameter values are given in supplementary tables 1 and 2. To estimate the mean time until adaptation by mutation, we used one centrally located patch of 99 demes and a mutation rate of *μ* = 10^*-5*^, and to estimate the mean time until adaptation by migration, we used two centrally located patches of 99 demes separated by a varying number of demes, with one patch initialized to have the *B* allele at frequency 0.8. In each case, we measured the “time to adaptation” as the amount of time until at least 100 *B* alleles were present in the previously unadapted patch. Figure 4 summarizes how the results compare to theory, excluding parameter combinations that violate the assumptions discussed above, or where a majority of simulations did not adapt by 25,000 generations. Supplementary figures 1 and 2 show all times, and depictions of typical simulation runs are shown in supplementary figures 3, 4, 5, 6, 7, 8, 9, and 10.

**Figure 4.**
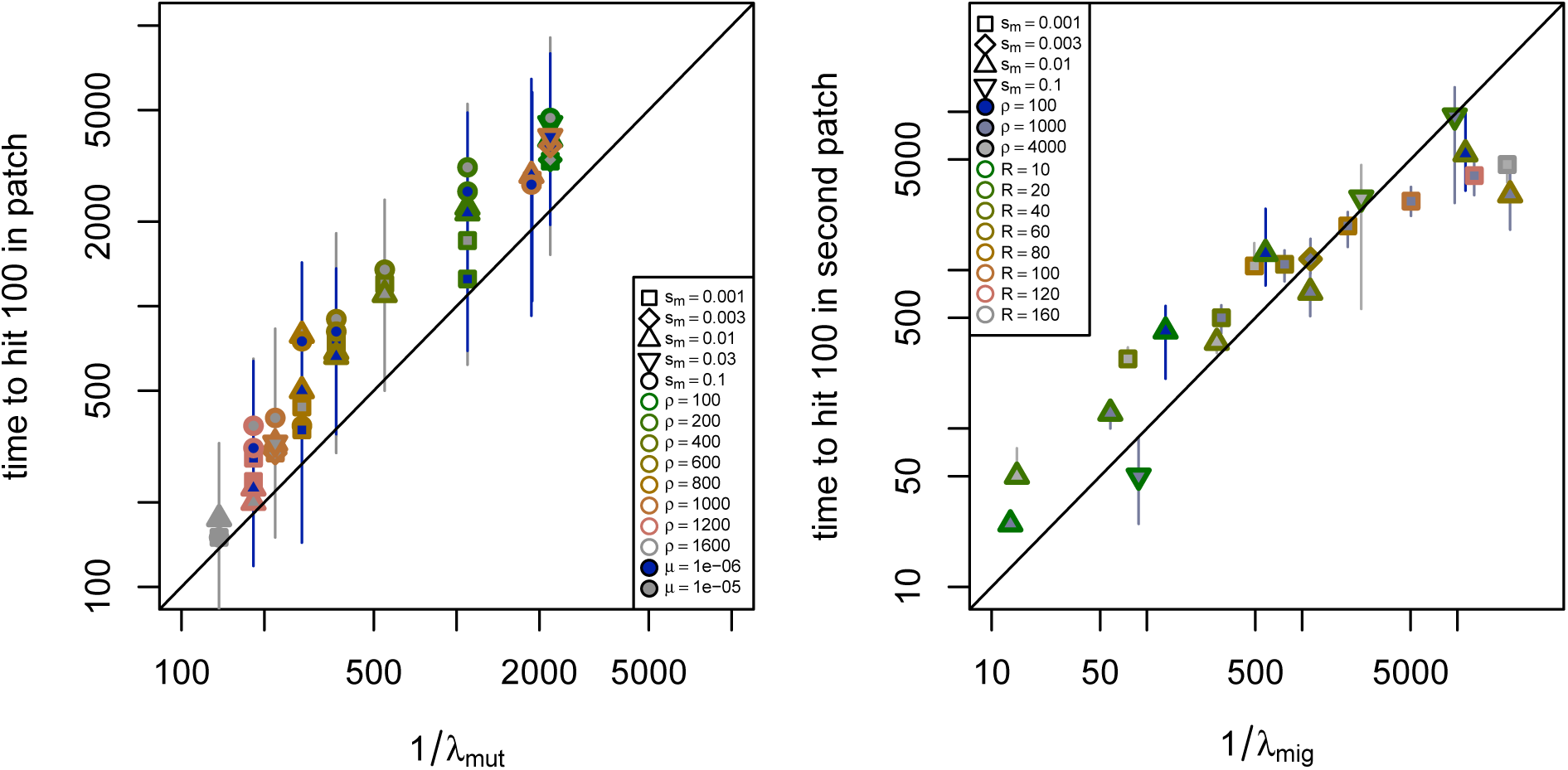
**(left)** Time until adaptation by mutation in an isolated patch for different combinations of selection coefficient on patches (*s*_*m*_), population density (*ρ*), mutation rate (*μ*), and distance between patches (*R*), as observed in simulations (vertical axis), and as predicted from equation (2) (horizontal axis). In all simulations, *s*_*p*_ = 0.01. **(right)** Time until adaptation by migration from an already occupied patch to a new patch, as observed in simulations (vertical axis), and as predicted from equation (11), with *C* = 5 (horizontal axis). For each, parameter combinations and numbers of simulations are in supplemental tables 1 and 2; shown are the median (points) and interquartile range (lines) of the time until the patch had at least 100 adapted individuals, for each unique parameter combination. Only parameter combinations to which the theory here is applicable were used (precisely, if 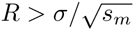, patch width larger than 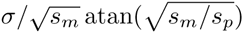 and *s*_*m*_ > 2σρ); the remaining combinations are shown in figures S1 and S2.

**Figure 5.**
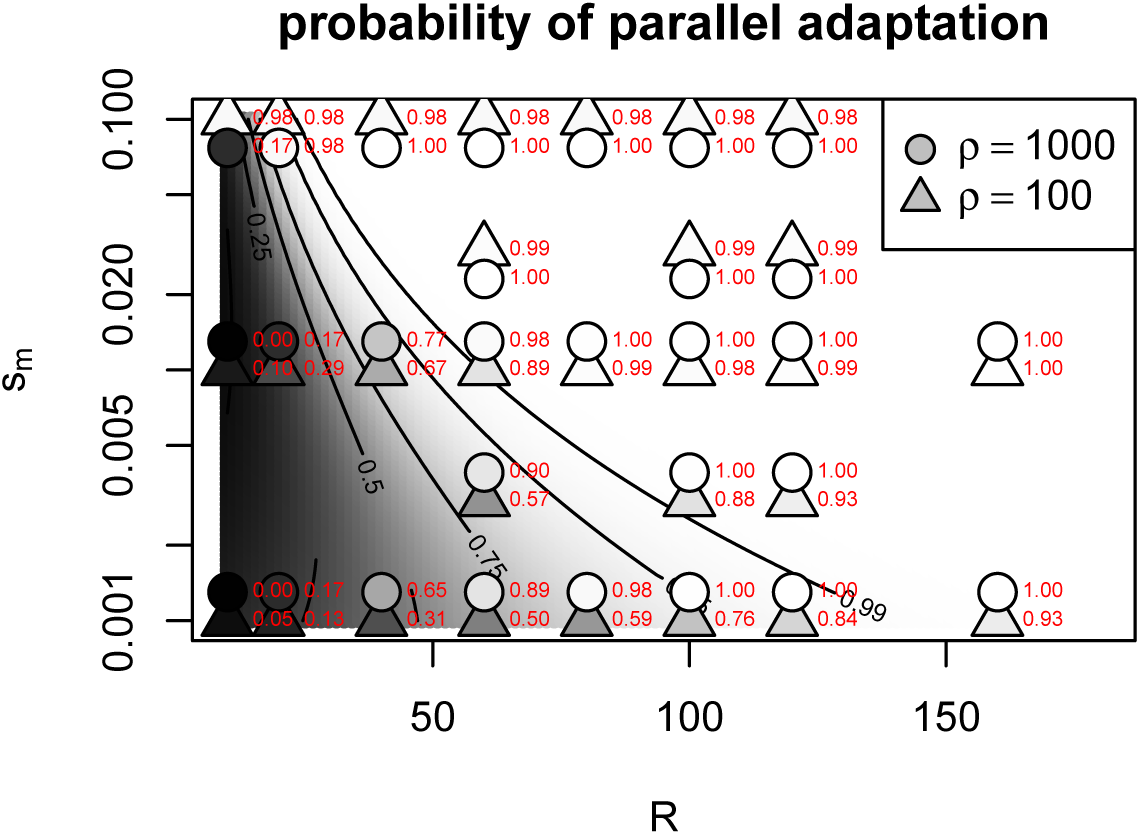
Probability that two patches evolve by independent mutations (i.e., convergently), as a function of their geographic separation (*R*) and the strength of selection between patches (*s*_*m*_). The background shades of grey and contours give the probability predicted by theory (equation (12), with *C* = 5). Points correspond to parameter combinations of the simulations in figure 4, with the shade of grey and the red numeric label indicating the same probability estimated from these simulations (see text for details). The two densities, *ρ* = 100 and *ρ* = 1000 would lie directly on top of each other, so the *ρ* = 1000 points have been shifted vertically.

The agreement is reasonable for both. The theoretical value 1*/λ*_mut_ underestimates the mean time to mutation by a factor of around 2 that increases slightly with *s*_*m*_. This is to be expected for two reasons: First, we compute the time to reach 100 *B* alleles, while theory predicts the time until the progenitor of those 100 *B* alleles arose. Second, the expression for *λ*_mut_ neglects effects near the boundary of the patch, and the larger *s*_*m*_ is, the harder it is for mutations that arise near the edge of the patch to establish. The theoretical values 1*/λ*_mig_ again has a margin of error of a factor of about 2. Supplementary figure 10 shows simulations at more parameter values.

### The probability of parallel adaptation between patches

We now turn to the question of whether the separated patches adapt by parallel genetic changes or by the spread of a migrant allele between the patches. As only a single mutation is required for individuals to adapt to the poor habitat patch, subsequent mutations that arise after an allele becomes established in the patch gain no selective benefit. Similarly, an allele introduced into a patch by migration will not be favored by selection to spread, if the patch has already been colonized. Therefore, mutations selectively exclude each other from patches, over short time scales, and will only slowly mix together over longer time scales by drift and migration, an assumption we also made in [Ralph and Coop, 2010]. In reality, different mutations will likely not be truly selectively equivalent, in which case this mixing occurs over a time-scale dictated by the difference in selection coefficients.

We assume that once a *B* allele is introduced (by migration or mutation) it becomes established in the poor habitat patch rapidly if it escapes loss by demographic stochasticity. Under this approximation, and the “selective exclusion” just discussed, after one of the patches becomes colonized by a single *B* allele, the other patch will become colonized by migration or mutation, but not by both. As such, the question of how the second patch adapts simply comes down to whether a new mutation or a migrant allele is the first to become established in the second patch. To work out how adaptation is likely to proceed, we can compare expressions (2) and (11) above for the effective rates of adaptation by new mutation and by migration. We work in one dimension, as the square root term appearing for two dimensions is relatively unimportant.

We first consider the order of magnitude that our parameters need to be on in order for adaptation via mutation or migration to dominate. Let *γ* = min(1*,s*_*m*_*/p*_*e*_) and *w* = *A/A*′ – since *A* is the area of the not-yet-adapted patch and *A*′ is the area of its closest 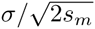 strip, *w* is approximately the width of the patch in units of 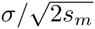. Effective migration and mutation rates will be on the same order if 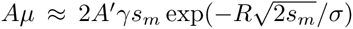, where *R* is the distance between the patches. In other words, the migrational analogue of “mutational influx” (*μρA*) 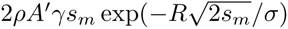, which depends fairly strongly on the selective disadvantage *s*_*m*_ between patches. Equivalently, the rates are roughly equal if 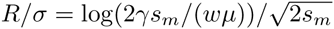, which gives the critical gap size past which adaptation will be mostly parallel in terms of selection coefficients, patch width, and mutation rate.

If we take *μ* = 10^−5^, the patch width to be ten times the length scale 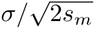 so *w* = 10 (and - log(*wμ*) ≈ 9.2), and *s*_*m*_ *> p*_*e*_ so that *γ* = 1, then adaptation is mostly parallel for patches separated by gaps larger than 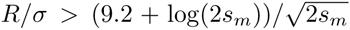. If the selective pressure is fairly strong (say, *s*_*m*_ = .05), then for convergence to be likely, the distance between patches must be at least 21.8 times the dispersal distance (i.e., *R/σ >* 21.8). If *s*_*m*_ is much smaller, say *s*_*m*_ = .001, the required distance increases to *R/σ >* 67.

If the mutation rate is higher – say, *μ* = 10^−3^ – the required separation between patches is only reduced to *R/σ >* 7.3 with *s*_*m*_ = .05. If *s*_*m*_ = .001, then with this higher mutational influx this calculation predicts that mutation will *always* be faster than migration – however, this should be taken with caution, since as discussed above, this model does not hold if *R* is of the same order as *σ* or if *s*_*m*_ is small enough that local drift is more important.

We can go beyond these rough calculations to find the probability of parallel adaptation if we are willing to take our approximations at face value. Doing so, the time until the first of the two patches is colonized by the *B* allele will be approximately exponentially distributed with mean 1*/*(2*λ*_mut_). Following this, the time until the second patch is subsequently colonized (via either migration or new mutation) will be exponentially distributed with mean 1*/*(*λ*_mut_ + *λ*_mig_). Both these rates scale linearly with population density (*ρ*) and the probability of establishment of a rare allele (*p*_*e*_ *≈* 2*s*_*b*_*/ξ*^2^, for *p*_*e*_ *> s*_*m*_), so that the overall time to adaptation is robustly predicted to increase with offspring variance (*ξ*^2^) and decrease with population density and beneficial selection coefficient. Furthermore, the probability that the second adaptation is a new mutation, i.e., the probability of parallel adaptation, with *γ* = min(1*,s*_*m*_*/p*_*e*_), is

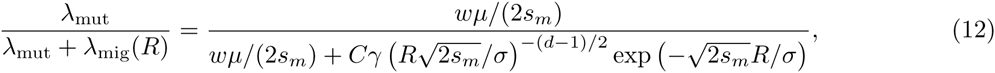

so that the probability of parallel mutation should increase approximately logistically with the distance between the patches, on the spatial scale 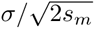.

We tested this using the same simulations as figure 4, by using the empirical distributions of the respective times to adaptation to estimate the probability, for each parameter combination, that a new, successful mutation appears before a successful migrant arrives from another, already adapted patch. The results are compared to (12) in figure 5.

### Multiple patches

Suppose that instead of patches there are many, each patch adapting through either migration or mutation. If the times between successive establishments are the result of many nearly-independent attempts with small probability of success, the process of occupation is well approximated by a Markov chain, whose state records the labeling of patches according to the colonizing allele (or none, for those that haven’t yet adapted).

If not yet adapted, a focal patch of area *A* will acquire the adaptation by new mutation at rate 2*s*_*p*_*μρA/ξ*^2^ (equation (2)). Suppose that patches 1*,…,k* are already adapted, and that these patches are at distances *R*_1_*,…,R*_*k*_ away from this unadapted patch. The total rate of adaptation through migration, from equation(11) if patches do not interfere with each other, is

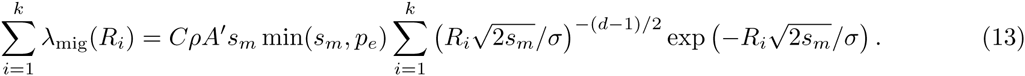

If the patch adapts through migration, the probability the adapted allele comes from patch *i* is

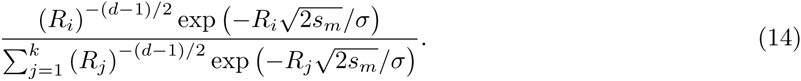

Since the compound parameter *s*_*p*_*ρ* is common to both migration and mutation rates, it functions only as a time scaling of the process, and therefore has no effect on the final configuration, namely, the sets of patches sharing the genetic basis of their adaptive response. We can also rescale time by the typical patch size, and introduce a parameter, say 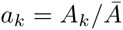, making the properties (other than the time scaling) of the discrete model independent of the *numerical sizes* of the demes themselves. This is complementary to the results of Pennings and Hermisson [2006], who showed that multiple mutations are likely to arise *within* a panmictic deme if the population-scaled mutation rate 2*Nμ* is greater than 1.

We can say more if we suppose that all patches have the same area and are the same distance from each other (i.e., the “island model”). The resulting process is a continuous-time version of the “Chinese restaurant process” described in Aldous [1985], and so the final partition of types across demes has the Ewens sampling distribution with parameter 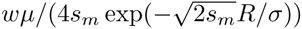. Now we can compute most properties we might want about the process. For instance, the proportion of demes that shares the same origin as a randomly sampled deme is approximately Beta distributed – see Donnelly and Joyce [1989]. The high connectedness of the discrete deme island model means that the expected number of distinct alleles grows with the log of the number of demes. This strongly contrasts with the continuous spatial model where the local nature of dispersal means that doubling the species range will double the number of mutations expected.

### Length of the hitchhiking haplotype

If a patch adapts through new mutation or a rare migrant lineage, the genomic region surrounding the selected site will hitchhike along with it [Maynard Smith and Haigh, 1974], so that at least initially, all adapted individuals within a patch share a fairly long stretch of haplotype. Pairs of adapted individuals within a patch will initially share a haplotype of mean genetic length of about log(*ρAs*_*p*_)*/s*_*p*_ around the selected site, as long as the patch is reasonably well mixed by dispersal (otherwise see Barton et al. [2013]). This association gets slowly whittled down by recombination during interbreeding at the edge of the patch, but there will always be longer LD nearby to the selected site [Barton, 1979].

When an already adapted patch colonizes another through migration, the newly colonized patch will inherit a long piece of haplotype around the selected site from the originating patch. The genetic length of this haplotype is roughly inversely proportional to the number of generations the allele spends “in transit”, because while very rare, the allele will be mostly in heterozygotes, and so each generation provides another opportunity for a different haplotype to recombine closer to the transiting *B* allele. A large, linked haplotype may still arrive and fix in the new patch, in which case the haplotype has literally hitchhiked across geography! Figure 6 shows a simulation of such an event, including the lineage that founds an adaptive population on a second patch, and the length of the haplotype shared between this lineage and one in the original patch. The time that the lineage is outside the region where the *B* allele is common (dark red in Figure 6), the haplotype that accompanies it is broken down rapidly. After the lineage establishes on the patch, the rate of decay of the haplotype is slowed significantly, since most others with which it recombines have similar haplotypic backgrounds.

**Figure 6.**
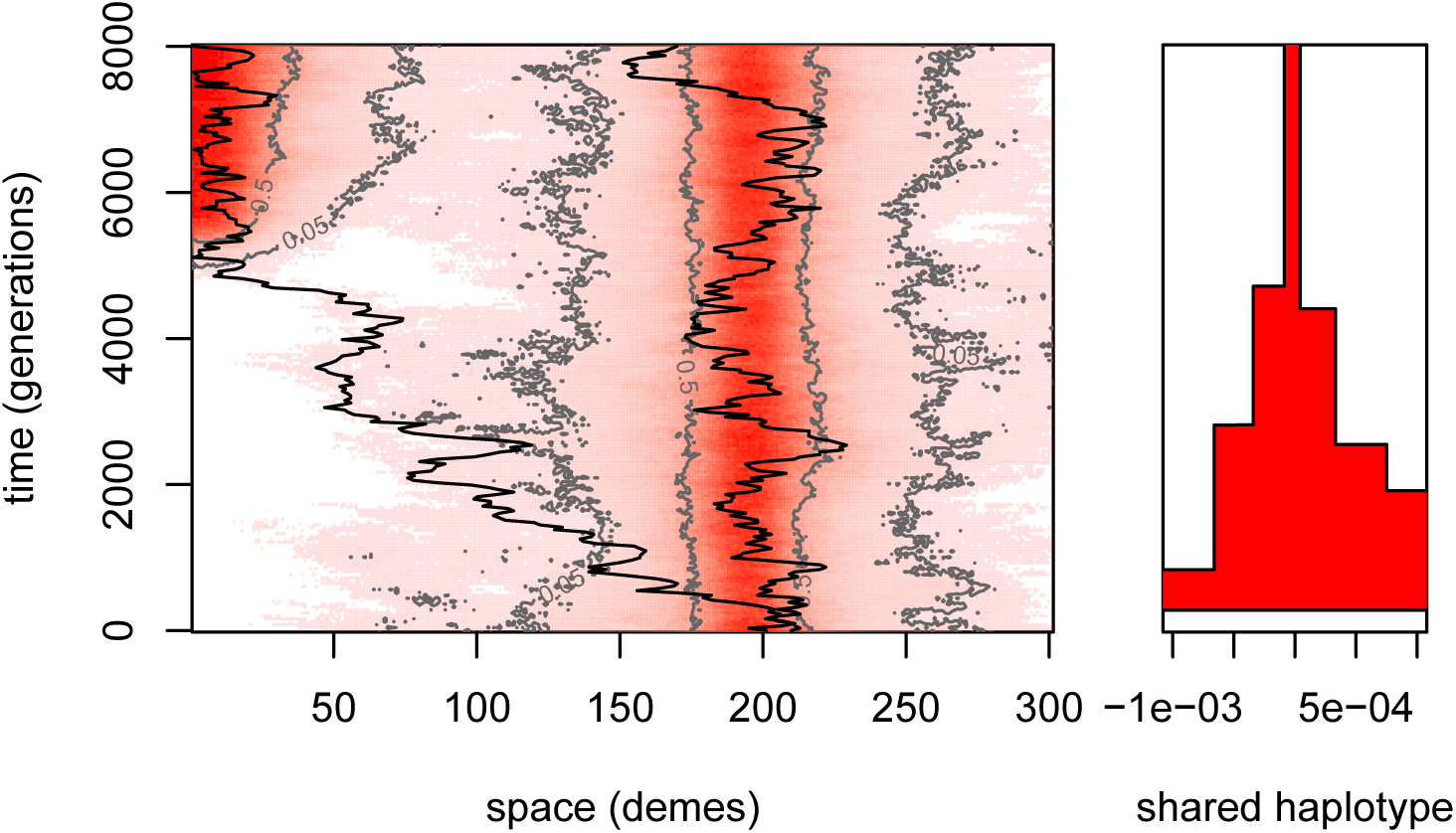
**Adaptation by migration:** An initially unadapted patch (10 demes at the left end of the range) is colonized around generation 5000; a representative lineage is shown in the new patch, which follows the original migrant “trunk” lineage between the patches and coalesces with a lineage in the unadapted patch. In the left panel, as in Figure 2, color and contours show allele frequency across space and time; other simulation parameters are as in Figures 1B and 2. The right panel shows (in red) the extent of the shared haplotype between the two lineages around the selected locus (at position 0), which decreases as time progresses.

Above we argued that a good model for this transit is a single Brownian trunk lineage surrounded by a cloud of close relatives of random size *K* whose chance of surviving until *t* generations is 1 − *k*_*e*_(*t*). Consider a single such family, and let *τ* be the (random) time at which it first hits the new patch, with *τ* = *∞* if it never does. We assume that the length of the hitchhiking segment depends only on the time spent by the trunk lineage of the first successful migrant “in transit” between the patches, i.e., *τ* conditioned on *τ < ∞*. In so doing, we neglect recombination between *B* alleles (since they are at low density in transit), and the possibility that more than one successful migrant family is in transit at once (so that faster migrants would be more likely to have arrived first).

Since each generation spent in transit provides an opportunity for recombination, if recombination is Poisson, the length of the haplotype (in Morgans) initially shared between populations on each side of the selected locus is exponentially distributed with mean *τ*. Therefore, if *L* is the length of hitchhiking segment on, say, the right of the selected locus, then

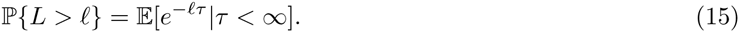

Computing this depends on 1 − *k*_*e*_(*t*), the probability a family survives until *t*. Since 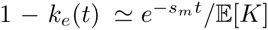, we approximate the lifetime distribution of a family by an exponential with mean 1*/s*_*m*_. We can then use standard results on hitting times of *d*-dimensional Brownian motion that is killed at rate *s*_*m*_ (see Borodin and Salminen [2002] 2.2.0.1 and 4.2.0.1). In particular, if the patch is circular with radius *w* and lies at distance *R* from the already adapted patch, then

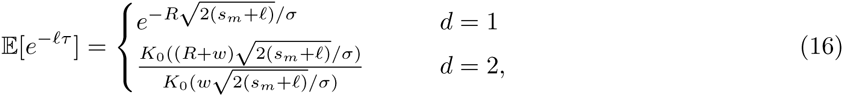

where *K*_0_ is a modified Bessel function of the second kind. We are interested in lineages that manage to reach the patch before being killed, i.e., having *τ < ∞*, which occurs with probability P{*τ < ∞*} = lim_*→*0_ 𝔼 [exp(- *lτ*)].

To keep the expressions simple, in the remainder of this section we only compute quantities for one dimension. By Bayes’ rule,

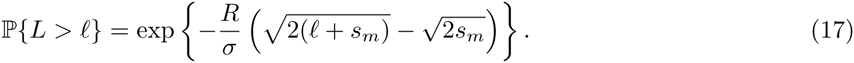

This can be differentiated to find that the expected transit time is

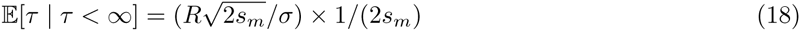

and that 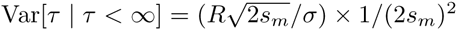. (A saddle point approximation provides an alternative route to these expressions.)

The form of equation (17) implies that if *Y* is an exponential random variable with rate 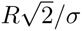, then *L* has the same distribution as 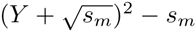. Furthermore, the expected length of shared hitchhiking haplotype is

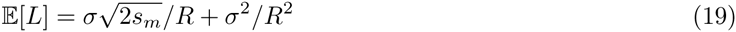

and 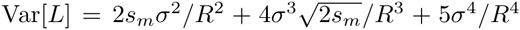. For two dimensions, asymptotics of Bessel functions show that *L* has the same tail behavior, but how other properties might change is unclear.

As a rule of thumb, equation (18) means that families who successfully establish move at speed 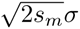 towards the new patch – if *s*_*m*_, the strength of the selection against them, is smaller, the need to move quickly is less imperative. Then, equation (19) means that the haplotype length is roughly the length one would expect given the mean transit time, so more weakly deleterious transiting alleles arrive with them shorter haplotypes. Note that we have become used to seeing *σ* divided by 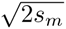, rather than multiplied; it appears here because 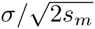 is a length scale, and to convert it to a speed this is divided by 1*/*2*s*_*m*_.

### Applications

Coat color in the rock pocket mouse *Chaetodipus intermedius* is a classic example of local adaptation [Benson, 1933, Dice and Blossom, 1937]. These mice live on rocky outcrops throughout the southwestern United States and northern Mexico, and generally have a light pelage similar to the predominant rock color. However, in some regions these mice live on darker substrates (e.g., old lava flows), and in these patches have much darker pigmentation, presumably to avoid visual predators. Some of the largest patches of dark rock range from 10km to 100km wide and lie 50–400km from each other, and dark-colored populations of *C. intermedius* have been found on many of these. (However, patches of all sizes occur across all scales in a heterogeneous manner.) Nachman et al. [2003] demonstrated that on one of these flows (Pinacate), much of the change in coloration is due to an allele at the MC1R locus. This dark allele differs from the light allele by four amino acid changes, and has a dominant or partially dominant effect depending on the measure of coat color. The Pinacate allele is not present in a number of other populations with dark pelage, suggesting these populations have adapted in parallel [Nachman et al., 2003, Hoekstra and Nachman, 2003]. However, Hoekstra et al. [2005] reasoned that, elsewhere in the range, multiple close dark outcrops may share a dark phenotype whose genetic basis has been spread by migration despite intervening light habitat.

A key parameter above was the dispersal distance divided by the square root of strength of selection against the focal allele between patches, 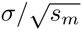. Hoekstra et al. [2004] studied the frequency of the dark MC1R allele and coat color phenotypes, at sites across the (dark) Pinacate lava flow and at two nearby sites in light-colored rock. On the lava flow the dark MC1R allele is at 86% frequency, while at a site with light substrate 12km to the west (Tule), the frequency is 3%. The dark allele was entirely absent from Christmas pass, a site with light substrate 7km north of Tule, and 3km further from the lava flow. In the other direction, the dark MC1R allele was at 34% at a site with light substrate 10km to the east of the flow (O’Neill). Note that such apparent asymmetry is expected, since as noted above the migration–selection equilibrium can be highly stochastic. These numbers give us a sense of a plausible range of the parameters. Assuming the dark allele is at 50% frequency at the edge of the lava flow, we can fit formula (3) to these frequencies (similar to Hoekstra et al. [2004]). Doing this for Tule we obtain 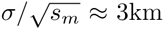, and for O’Neill, 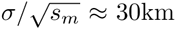, giving us a range of cline widths. We also need estimate of *s*_*m*_. Using *σ ≈* 1km [French et al., 1968, Allred and Beck, 1963], these cline widths imply that *s*_*m*_ = 1*/*9 and 1*/*900 are reasonable values.

The mutational target size *μ* for the trait is unclear. While the Pinacate dark haplotype differs from the light haplotype at four amino acid residues, it is likely that not all of these changes are needed for a population to begin to adapt. Also, there a number of genes besides MC1R at which adaptive changes affecting pigmentation have been identified in closely related species and more broadly across vertebrates [Hoekstra, 2006]. To span a range of plausible values, we use a low mutation rate of *μ* = 10^−8^ (a single base pair), and a high mutation rate *μ* = 10^−5^ (a kilobase). Finally, we set *A* = 100km^2^ (roughly the size of the Pinacate patch). In Figure 7 we show the dependence of the probability of parallel mutation on the distance between lava flow patches using these parameters, showing that parallel mutation should become likely over a scale of tens to a few hundred kilometers between patches.

**Figure 7.**
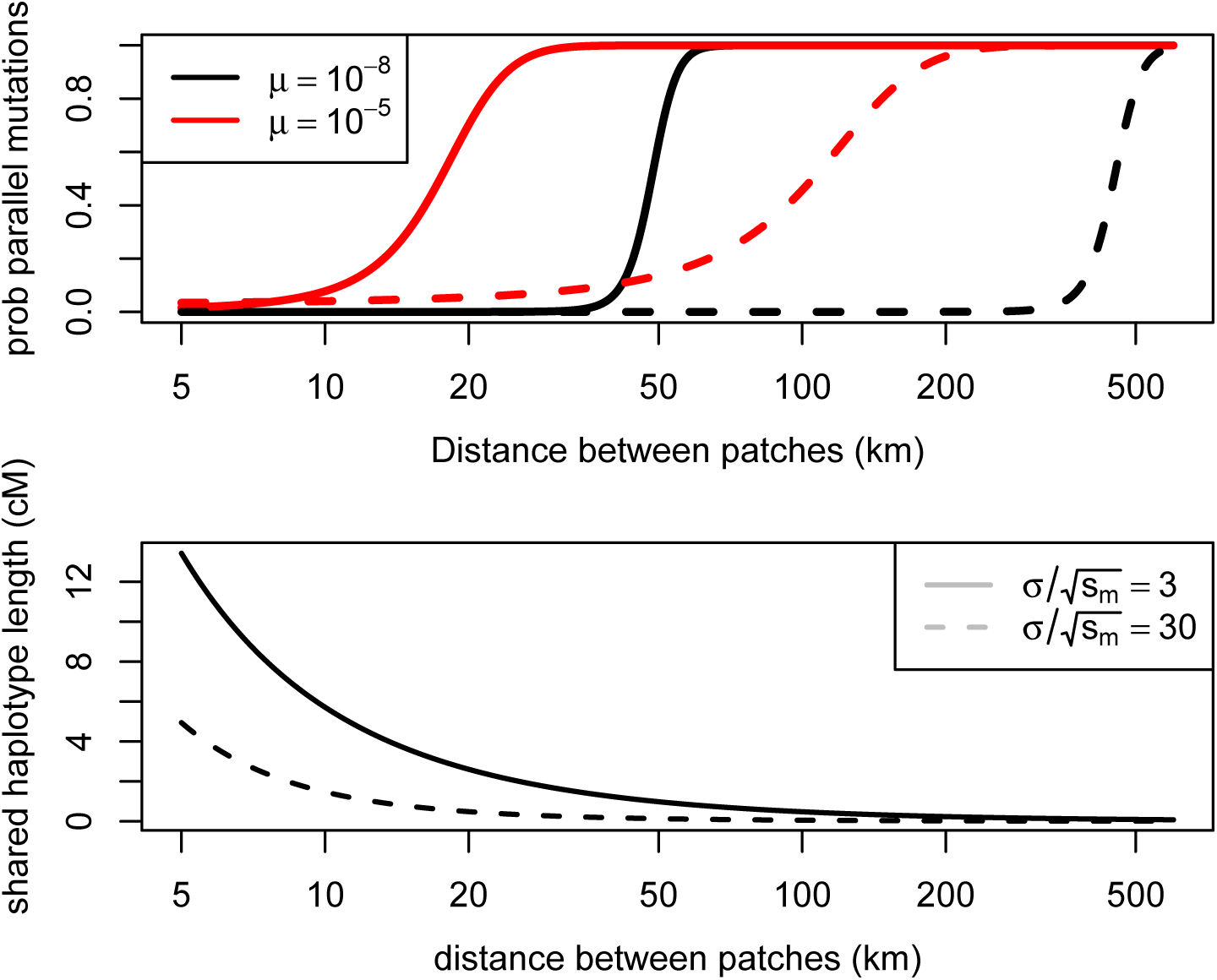
Top panel: The probability of parallel mutation as a function of the distance between environmental patches for two different cline widths and two different mutation rates (using equation (12)). The parameters were chosen to match the example of *Chaetodipus intermedius*; see text. Bottom panel: The initial genetic length of the haplotype shared between patches due to adaptation via migration as a function of the distance between environmental patches. As a rough rule of thumb 1cM is approximately 10^6^ bases in a number of mammalian species.

**Figure 8.**
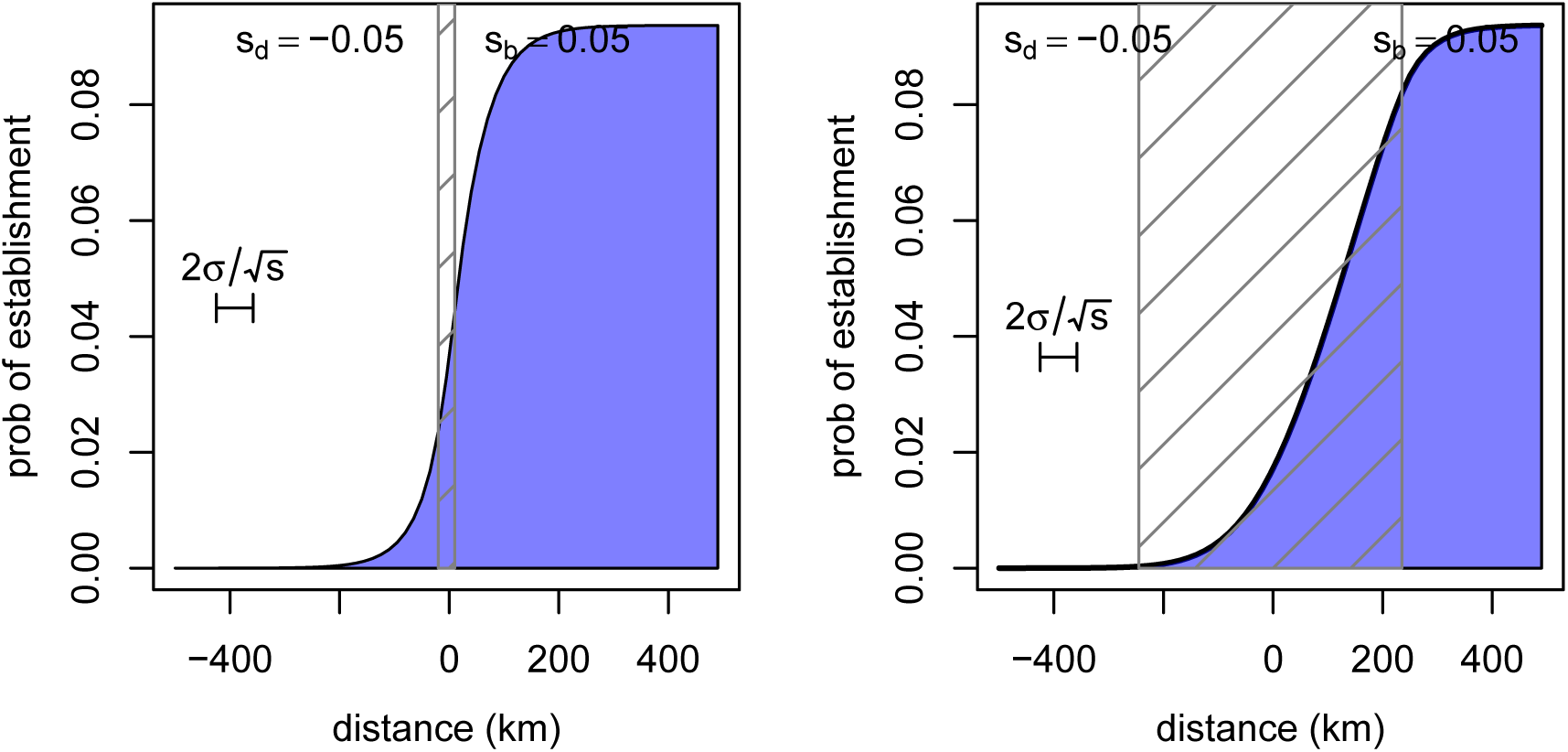
The probability of establishment of a new mutant allele as a function of distance from the edge of the region where it is beneficial, in an abrupt transition (left) and a gradual transition (right). The allele is deleterious to the left and beneficial to the right (with selection coefficients *±*0.05 respectively); and the selection coefficient interpolates linearly between these in the central hatched region. The number of offspring is Poisson. The underlying landscape has demes every 15km and a migration rate of 0.5 between neighboring demes; the probability was found by numerically solving the equation for the probability of establishment of a multi-type branching process. Note that the approximation used in the text is here *p*_*e*_ *≈* 2*s*_*p*_*/ξ*^2^ = 0.1, agreeing with the value in the center of the patch.

Given the large selection coefficient associated with the dark allele on the dark substrate, we expect the initial haplotype associated with either a new mutation or migrant allele to be large. Figure 7 also shows how long the founding haplotype shared between populations is expected to be, from equation (19). The initial length can be quite long between geographically close patches (tens of kilometers). However, for the wider cline width 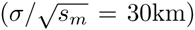, adaptation by migration can still be likely for patches 100km apart, but the shared basis may be hard to detect, as the length of shared haplotype can be quite short.

## Discussion

This paper is an investigation into the basic question: What is the spatial resolution of convergent local adaptation? In other words, over what spatial scale of environmental patchiness will the process of adaptation develop independent solutions to evolutionary problems? The answer to this depends most strongly on 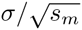, the dispersal distance divided by the square root of the strength of selection against the allele between patches. It depends much more weakly on the selective benefit within the patches or (perhaps surprisingly) the population density, although these two factors will determine the time-scale over which adaptation will occur (and note that population density could affect the selection coefficients). This is in contrast with models of panmictic populations [Hermisson and Pennings, 2005, Messer and Petrov, 2013, Wilson et al., 2014] and geographically spread populations adapting to homogeneous selection pressures [Ralph and Coop, 2010], where the probability of multiple, independently arising adaptive alleles increases with the population size. However, in all of these models the dependence on the beneficial selection coefficient is absent or weak, due to the fact that selection both aids establishment of new alleles and the spread of existing alleles (but see Wilson et al. [2014] for the complications of varying population sizes).

We have also shown that while weaker selection against alleles will make sharing of adaptations between patches easier, it also makes it harder to spot such sharing, since the lucky alleles that manage to colonize new patches move slower, and thus carry a shorter shared haplotype. This issue is amplified by the fact that the length of haplotype shared within patches decays over time, potentially making the identification of shared adaptations to old selection pressures difficult.

Perhaps the most useful rule-of-thumb quantities we found were the following. The effective rate of migration into an as-yet-unadapted patch from an already-adapted patch distance *R* away – the analogue of the mutational influx 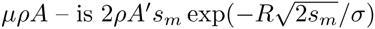. Equivalently, the critical gap size between patches past which adaptation is likely independent is 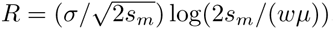. Finally, successfully transiting migrant lineages move at rate 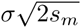, and so shared haplotype lengths between patches will be of order 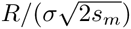.

In developing the set of approximations in the paper we have ignored a number of complicating factors. We now briefly discuss these.

### Standing variation

We have focused on relative rates of adaptation, since in applications where adaptation has occurred, the question is whether adaptations in distinct patches have appeared independently or not. However, any adaptation that does occur may have to make use of standing variation, if mutation rates are low. The case of a panmictic population was studied by Hermisson and Pennings [2005], and we study the case of a continuous, spatial population in Ralph and Coop [2015]. If parallelism in local adaptation of the sort we study here is due to standing variation rather than new mutation, then the dynamics of adaptation should not depend strongly on migration patterns (but the initial spatial distribution of standing variation may).

### Dominance

We have mostly ignored the issue of dominance by dealing with essentially haploid models, and appealing to the fact that the dynamics we study occur where the mutation is rare, and hence mostly present only in heterozygotes. Our results should hold as a good approximation to dominant and partially dominant alleles (with *s*_*m*_ the selection against heterozygotes). If, however, the mutation is recessive, then it is essentially neutral where rare, and so would encounter much less resistance to spreading between patches. The shape of the cline obtained is given by Haldane [1948]. This is counteracted, however, by the increased difficulty with which the mutation would establish once arriving in a new patch, if the beneficial effect is also recessive. As such it is not clear what our intuition should be about the contribution of recessive alleles to adaptation via migration. Further work is needed to put empirical observations of local adaptation by recessive alleles in a theoretical context. It is similarly unclear how the model should extend to a polygenic trait.

### Decay of shared haplotypes

To provide context for the results on shared haplotype length in section *Haplotypes Shared Between Patches* it is important to also understand the process by which haplotypes are whittled down within patches. The initial haplotype that sweeps within a patch will be dispersed over time by recombination. Likewise, the haplotype that is shared between patches coadapted by migration will also break down (equation 19). However, a long time after the initial sweep, we may still expect to find individuals within the patch sharing longer haplotypes around the selected locus than with individuals elsewhere, since selection against migrants decreases mean coalescence times within the patch near the selected locus. The literature on clines (e.g., Barton [1979]) has important information, but more work is needed to provide robust estimates for these processes. Questions about the genomic length-scale of signals of sweeps shared by migration have also been addressed in discrete population settings [Slatkin and Wiehe, 1998, Kim and Maruki, 2011], reviewed in Barton [2000]. This work has shown that the length of the shared swept haplotype is often significantly shorter than the sweep within each patch, resulting in a pattern of shoulders of elevated *F*_*ST*_ between adapted populations some distance away from the shared selected allele. It would be of interest to see how similar patterns can arise in a continuous population setting, as a way of uniting these results.

### Long distance migration

We have also ignored the possibility of very long distance migration, instead focusing on local dispersal (hence Gaussian by the central limit theorem). However, dispersal distributions can be very heavy tailed, with a small fraction of individuals moving very long distances indeed [Levin et al., 2003, Reynolds and Rhodes, 2009]. In addition, over long time-scales, very rare chance events (mice carried off by hurricanes and the like; Censky et al. [1998], Nathan et al. [2008]) could play a role in spreading migrant alleles if adaptation by other means is sufficiently unlikely. Such tail events could greatly increase the probability of shared adaptation above that predicted by our model. Furthermore, if adaptive alleles do move between distant patches via rare, long distance migration then they will be associated with a much longer shared haplotype than predicted by equation (19). As such, we view our results as a null model by which the contribution of long distribution migrants to adaptation could be empirically judged.

### Other geographic models

We have studied circular patches of habitat at long distances from each other. Real habitat geometry can be much more complex, e.g., with archipelagos of patches of varying sizes, or patches connected by long, skinny corridors, for instance. The work of Cantrell and Cosner [1991] comes closest to a general theory of balanced polymorphisms in such habitats. It is possible that our techniques could be applied in their much more general setting, as both are based, fundamentally, on branching process approximations. It is also interesting to think about the probability of convergent adaptation to continuously varying environments, e.g. replicated environmental clines.

### Concluding thoughts

The falling cost of population genomic sequencing means that we will soon have the opportunity to study the interplay of adaptation with geography and ecology across many populations within a species. Our work suggests that even quite geographically close populations may be forced to locally adapt by repeated, convergent, *de novo* mutation when migration is geographically limited and selective pressures are divergent. Thus, systems where populations have been repeatedly subject to strong local selection pressures may offer the opportunity to study highly replicated convergent adaptation within a similar genetic background [Stern, 2013]. Such empirical work will also strongly inform our understanding of the ability of gene flow to keep ecologically similar populations evolving in concert [Sexton et al., 2013]. Our results suggest that adaptation to shared environments is certainly no guarantee of a shared genetic basis to adaptation, suggesting that rapid adaptation to a shared environment could potentially drive speciation if the alleles that spread in each population fail to work well together [Kondrashov, 2003].

## Materials & Methods

### Simulation methods

First we briefly describe the simulations we used for illustration and validation (the R code used is provided as a supplement). We simulated forward-time dynamics of the number of alleles of each type in a rectangular grid (either one- or two-dimensional) of demes with fixed size *N*. Each generation, each individual independently chose to reproduce or not with a probability *r* depending on her type and location in the grid; locally beneficial alleles were more likely to reproduce. Each extant individual then either remained in the same location with probability 1 − *m* or else migrated a random number of steps in a uniformly chosen cardinal direction; for most simulations *m* = 0.2 and the probability of migrating *k* steps was proportional to 2^−^*k* for 1 *≤ k ≤* 5. In 2D, diagonal steps were also used. This gave us values of *σ* = 0.95 deme spacings in 1D and *σ* = 0.74 in 2D. Once migrants were distributed, each deme was uniformly resampled back down to *N* individuals. (Although we described the simulation in terms of individuals, we kept track only of total numbers in an equivalent way.)

The base probability of reproduction in each generation in simulations for type *b* alleles was *r* = 0.3; this was then multiplied by 1 + *s* to get the probability of reproduction for type *B*, where the value of *s* is either *s*_*m*_ or *s*_*p*_ depending on the individual’s location. This determines the values of *s*_*m*_ and *s*_*p*_ reported in the figures, and do not depend on the basic rate of reproduction. However, to obtain values for *s*_*p*_ and *s*_*m*_ when comparing theory to simulation, we computed the rate of intrinsic growth, i.e., the *s* so that the numbers of *B* alleles when rare would change by *e*^*st*^ after *t* generations in the absence of migration. (The resulting values are close to the first notion of *s*, but give better agreement with theory, which uses the second definition.)

To sample lineages, we first simulated the population dynamics forwards in time, then sampled lineages back through time by, in each generation, moving each lineage to a new deme with probability proportional to the reverse migration probability weighted by the number of *B* alleles in that deme in the previous time step. If more than one lineage was found in a deme with *n* alleles of type *B*, then each lineage picked a label uniformly from 1 *…n*, and those picking the same label coalesced. Since reproduction is Poisson, this correctly samples from the distribution of lineages given the population dynamics.

### Numerical calculation of the probability of establishment

When rare, copies of a new mutant allele are approximately independent and experience a uniform selective benefit; and can therefore be treated as a branching process. Furthermore, whether or not a new, beneficial mutation establishes or is lost to demographic stochasticity is determined by this initial phase where it is rare. Fortunately, the probability that a branching process dies out can be found as a fixed point of the generating function of the process [Jagers, 1975]. Therefore, we calculated explicitly the generating function for a spatial branching process with nearest-neighbor migration on a one-dimensional lattice and a Poisson number of offspring with mean 1+*s*, where *s* could vary by location, and iterated this forward to convergence to obtain 1 − *p*(*x*), the probability a single mutation appearing at *x* would fail to establish. We considered two situations: where *s* is a step function, and where it has a linear transition. These solutions are shown, and parameters described, in figure 8.

Here (and at other parameter choices) we see that the probability of establishment *p*(*x*) goes to the equilibrium value (approximately *p*_*e*_ = 2*s/ξ*^2^) within the patch; the transition is fairly symmetrical about the edge of the patch, even if the edge of the patch is not sharp. Additional experimentation indicated that the fit remains equally good for other parameter values, even if migration can move further than one deme and offspring numbers are not Poisson. This lends credence to our approximation that the integral of *p*(*x*) over the entire range is close to *p*_*e*_ multiplied by the area of the patch.

### The equilibrium frequency

For completeness, and clarity as to the scalings on the relevant parameters, here we provide a derivation of the differential equations referred to above equation (3), and establish the asymptotics given in that equation. One route to the “equilibrium frequency” of the allele outside the range where it is advantageous is as follows; see Slatkin [1973] or Barton [1987] (or Kolmogorov et al. [1991] or Fisher [1937] or Haldane [1948]) for other arguments in equivalent models, and see Etheridge [2000] and/or Dawson [1993] for a general framework for the stochastic processes below.

Suppose that the population is composed of a finite number of small demes of equal size *N* arranged in a regular grid, and that selection (for or against) the allele is given by the function *s*(*x*), with *x* denoting the spatial location. Each individual at location *x* reproduces at random, exponentially distributed intervals, producing a random number of offspring with distribution given by *X* who then all migrate to a new location chosen randomly from the distribution given by *x* + *R*, where they replace randomly chosen individuals. If *x* + *R* is outside of the range, then they perish. Each individual’s time until reproduction is exponentially distributed: the reproduction rate is 1 if it carries the original allele, or is 1 + *s*(*x*) if it carries the mutant allele. Suppose that the number of offspring *X* has mean *μ*; the variance of *X* will not enter into the formula (but assume *X* is well-behaved). Also suppose that the migration displacement *R* has mean zero and variance *σ*^2^; in more than one dimension, we mean that the components of the dispersal distance are uncorrelated and each have variance *σ*^2^.

Let 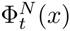 be the proportion of mutant alleles present at location *x* at time *t*, and Φ_*t*_(*x*) the process obtained by taking *N → ∞* (which we assume exists). Denote by *δ*_*x*_ a single unit at location *x*, so that e.g. 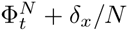 is the configuration after a mutant allele has been added to location *x*. For 0 *≤ ϕ ≤* 1, we also denote by 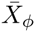 the random number of mutant alleles added if *X* new offspring carrying mutant alleles replace randomly chosen individuals in a deme where the mutant allele is at frequency *ϕ* (i.e. hypergeometric with parameters (*X, Nϕ, N*(1 − *ϕ*))); similarly, 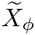 is the number lost if the new offspring do not carry the allele (i.e. hypergeometric with parameters (*X, N*(1 − *ϕ*)*,Nϕ*)). (We like to think of 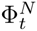 as a measure, but it does not hurt to think of Φ^*N*^ as a vector; we aren’t providing the rigorous justification here.) Then the above description implies that for any sufficiently nice function *f*(Φ) that

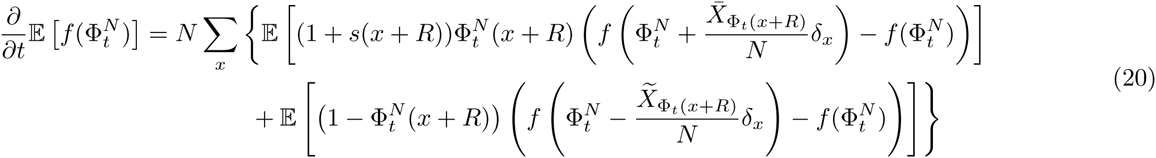

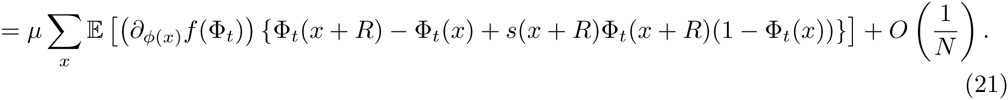

In the final expectation, *R* and Φ are independent. This follows by taking first-order terms in 1*/N* in the Taylor series for *f*, and the fact that 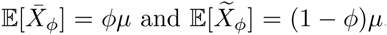. We can see two things from this: First, since this is a first-order differential operator, the limiting stochastic process Φ obtained as *N → ∞* is in fact deterministic (check by applying to *f*(Φ) = Φ(*x*)^2^ to find the variance). Second, if we want to rescale space as well to get the usual differential equation, we need to choose Var[*R*] = *σ*^2^ and *s*(*x*) to be of the same, small, order; this is another way of seeing that 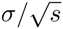 is the relevant length scale (as noted by Slatkin [1973]). More concretely, suppose that the grid size is ϵ *→* 0, that Var[*R*] = (*σ*ϵ)^2^, and that the strength of selection is *s*(*x*)ϵ, suppose that Φ_*t*_(*x*) is deterministic and twice differentiable, and let *ξ*(*t, x*) = Φ_*t/ϵ*_ (*x*); then the previous equation with *f*(Φ) = Φ(*x*) converges to the familiar form:

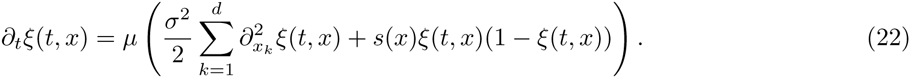

Here we have taken first the population size *N → ∞* and then the grid size ϵ *→* 0; we could alternatively take both limits together, but not if goes to zero too much faster than *N* grows. One reason for this is that at finite *N*, the process Φ_*t*_ is an irreducible finite-state Markov chain with absorbing states at 0 and 1; therefore, the inevitable outcome is extinction of one type or another, which is not the regime we want to study.

In one dimension, we are done (and discuss exact solutions in the Supporting Information); in higher dimensions, we are more interested in the mean frequency at a given distance *r* from a patch. If we take a radially symmetric patch centered at the origin (so *s* only depends on *r*), and let *ξ*(*t, r*) denote the mean occupation frequency at distance *r*, then the polar form of the Laplacian in *d* dimensions gives us that (22) is

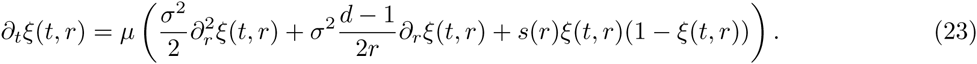

### Asymptotic solution for the equilibrium frequency

A radially symmetric equilibrium frequency *ξ*(*t, x*) = *q*(*|x|*), with *s*(*r*) = *−s <* 0 for all *r > r*_0_, solves for *r*_0_ *< r < ∞*,

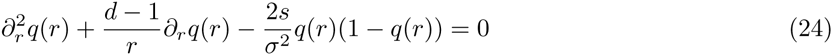

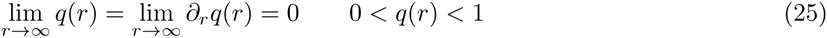

Since *q*(*r*) *→* 0 as *r → ∞*, so *q*(*r*)(1 − *q*(*r*)) *≈ q*(*r*), it can be shown that the true equilibrium frequency *q* is close, for large *r*, to the solution to

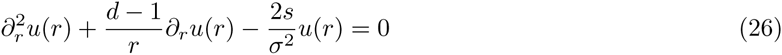

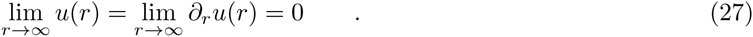

This has general solution given by a modified Bessel function: using Gradshteyn and Ryzhik [2007] 8.494.9, the general solution is

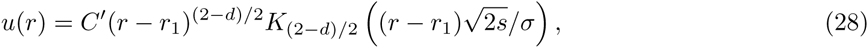

where *C*′ and *r*_1_ are chosen to match boundary conditions. Asymptotics of Bessel functions (Gradshteyn and Ryzhik [2007], 8.451.6) then imply that

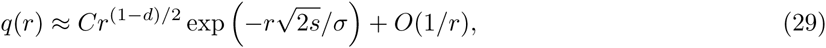

where *C* is a different constant.

### Hitting and occupation

Here we make a more precise argument to back up expression (11) for the migration rate. The argument made above in section *Heuristics* applies to general migration mechanisms, since it relies only on a decomposition of the migrant families upon hitting the new patch; but it is also imprecise in subtle ways that are difficult to formalize. Here we take a somewhat different tack, supposing that it suffices to model the spatial movement of a migrant family by following only the motion of the “trunk” (i.e., the red line in figure 3), and supposing this motion is Brownian, with variance *σ*^2^ per generation. (Recall *σ* is the dispersal distance.) We then use facts about Brownian motion and branching processes, to compute more precise versions of equations (4) and (5).

We are approximating the dynamics of the focal allele in the region further away than *r*_0_ from the patch as the sum of independent migrant families, each of whose dynamics are given by a spatial branching process (as depicted in Figure 3). Call *B*(*r*_0_) the region closer than *r*_0_ to the patch, and *∂B*(*r*_0_) its boundary. Denote by *γ*(*x*) the mean rate of outflux of migrant families from a point *x ∈ ∂B*(*r*_0_), i.e., the time-averaged density of individuals near a point *x* in *∂B*(*r*_0_) that are the founders of new migrant families. Expressions (4) and (5) are a simple product of the “outflux of families”, when in fact they should be an integral of *γ*(*x*) over possible locations. However, it will turn out that the integrand of equation (5) is well-approximated by a constant multiple of the integrand of (4).

First consider equation (4) for the equilibrium frequency. Suppose that *Z* is one such spatial branching process as above in section *The genealogy of migrant families*, started at time 0 with a single individual at *x*, and write *S* for the new patch. The *mean occupation measure* of *Z* in the region *S*, which we denote *u*(*x, S*), can be thought of informally as the expected total number of offspring of a family beginning at *x* that ever live in *S*. Let 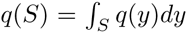 denote the total equilibrium frequency in *S*. This is decomposed in expression (4) as the sum of mean occupation densities of a constant outflux of branching processes from *∂B*(*r*_0_):

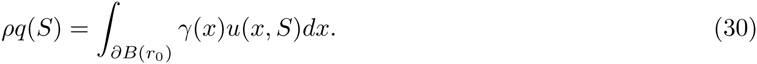

Now we will decompose *u*(*x, S*) under the assumption that the marginal distribution of the spatial motion of a single lineage is Brownian. Let *B*_*t*_ be a Brownian motion with variance *σ*^2^, and *τ*_*†*_ an independent Exponential(*s*_*m*_) time. The mean occupation time of *Z* spent in a region *S* is,

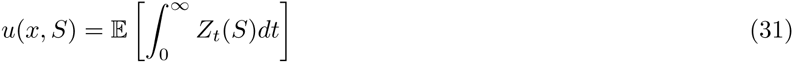

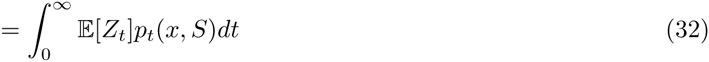

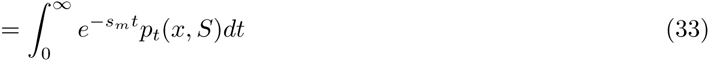

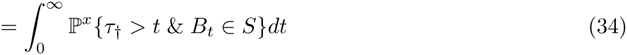

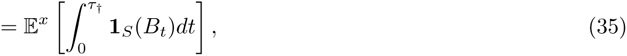

where ℙ^*x*^ gives probabilities for Brownian motion begun at *x* (i.e., *B*_0_ = *x*), and likewise 𝔼^*x*^. If we define *τ*_*S*_ to be the hitting time of *S* by the Brownian motion *B*, and *μ*_*S*_(*x*) to be the hitting distribution of *∂S* by *B*_*τS*_ conditioned on *τ*_*S*_ *< τ*_*†*_, by the strong Markov property this is equal to

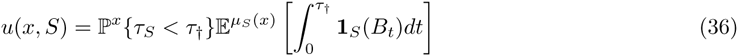

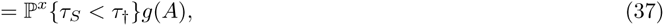

where now 𝔼^*μ*^ denotes expectations for Brownian motion for which the distribution of *B*_0_ is *μ*, and *g*(*A*) is defined to be the latter expectation, which does not depend on *x* if *S* is circular (with area *A*).

This form we can now compare to the expression (5) for the outflux of successful migrants. Consider the probability that a migrant family beginning with a single individual at *x* will ever establish in the new patch. It would be possible to analyze this probability directly, as in Barton [1987]; but here we take a simpler route, approximating this by the chance that the trunk hits the new patch, multiplied by the chance that at least one member of the family escapes demographic stochasticity and successfully establishes in the new patch. Write *h*(*x, S*) for the probability that the Brownian trunk hits the patch *S* before the family dies out, and *f*(*S*) for the chance that the family manages to establish in the new patch, given that it successfully arrives. (This is approximately independent of *x*.) As for *q*(*S*) above, expression (5) is properly an integral against the outflux *γ*(*x*):

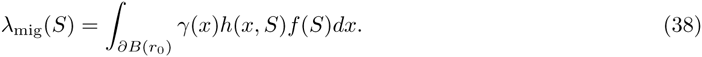

If we make the approximation that *Z* hits the new patch only if the trunk of *Z* does, and recall that 1 − *k*_*e*_(*t*) is the chance that *Z* survives for *t* generations, then

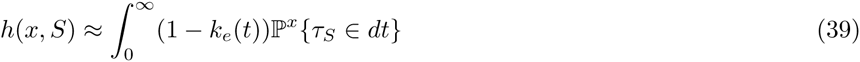

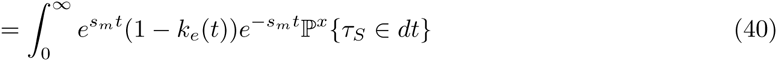

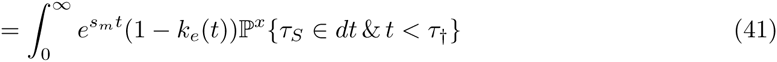

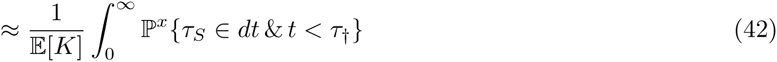

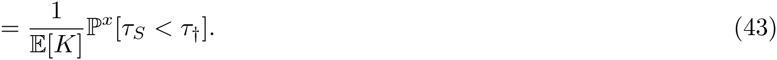

Therefore, we have that

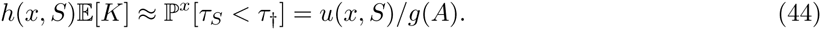

Since the integrands in expressions (30) and (38) only differ by a factor that (at least asymptotically) does not depend on the distance between *x* and *S*, we can obtain the migration rate by multiplying the equilibrium frequency by this factor. This factor will not depend on *A*, because although *f*(*A*) and *g*(*A*) in principle depend on the patch size (and geometry), the dependence is very weak. For instance, the width of a circular patch only very weakly affects the chance of establishment of a new migrant that appears on its edge, as long as the patch is large enough.

By equations (30) and (38),

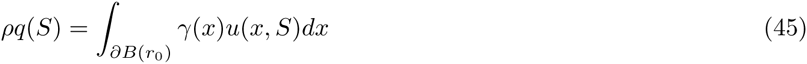

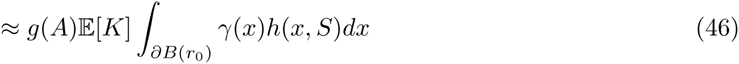

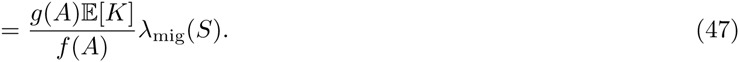

Once we show that *g*(*A*) *≈* 1*/*(2*s*_*m*_), we will have arrived at the result, equation (10).

The function *g*(*A*) is the expected amount of time that a Brownian motion begun on the edge of a disk of area *A* is expected to spend inside the disk before *τ*_*†*_. This is integral of the Green function for the Bessel process of the appropriate order, so using Borodin and Salminen [2002], and letting *w* be the width of the patch, in *d* = 1,

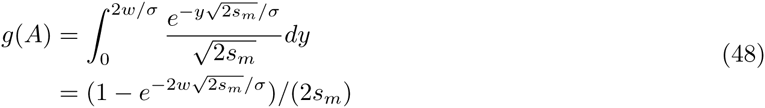

and in *d* = 2, by Gradshteyn and Ryzhik [2007] 5.56.2,

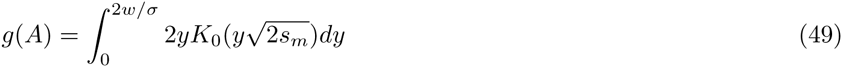

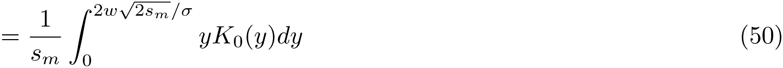

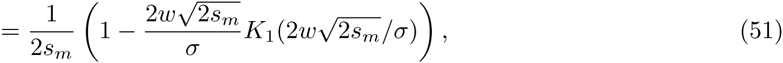

where *K*_0_ and *K* _1_ are modified Bessel functions of the second kind. In either case, *g*(*A*) *≈* 1*/*(2*s*_*m*_), which is the approximation we use in the main text.

### The integral *q*(*S*)

In the development above we need to approximate the integral of equation (3) across the area occupied by the new patch, 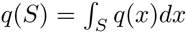. The precise answer depends on the shape and orientation of the patches; but we aim for a usable approximation, primarily in terms of the shortest distance from the old patch to the new patch, denoted *R*, and assuming *R* is large enough we can take equation (3) as an equality.

Since *q*(*R*) decreases as *|x|* increases,

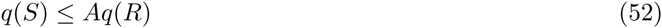

where *A* is the area of *S*. This will be a good approximation if *S* is small.

In one dimension, if *S* has length *l*,

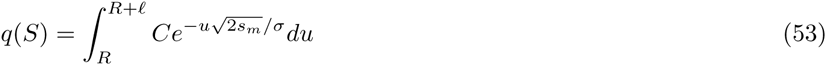

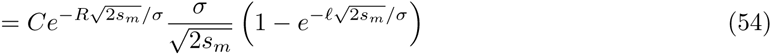

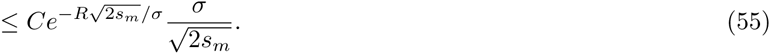

In two dimensions, suppose that *T* is a rectangle enclosing *S* with one axis aligned towards the original patch and transverse width *w*. Then

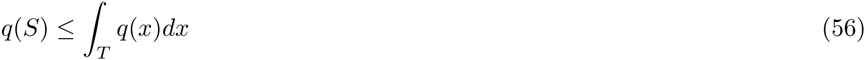

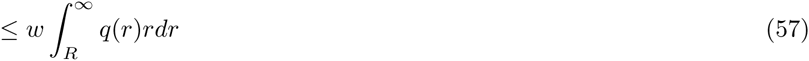

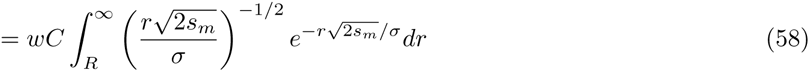

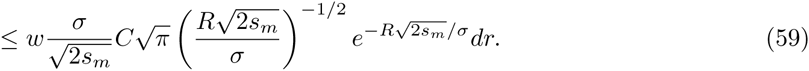

If we absorb 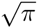 into the constant *C*, this is of the form 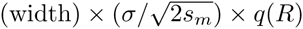, i.e., roughly the area, after replacing the length by 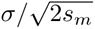.

In both cases, the upper bound is *q*(*S*) *≤ A*′ *q*(*x*), where *A*′ is the area of the parts of *S* that are no more than 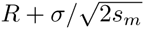 away from the original patch. Lower bounds could be obtained along similar lines.

## Acknowledgements

We would like to heartily thank Joachim Hermisson, Richard Neher, and an anonymous reviewer for a number of insightful suggestions, and catching several errors present in earlier drafts. We would also like to acknowledge Michael Nachman, Steve Evans, and Yaniv Brandvain for useful discussions, as well as Gideon Bradburd, and Simon Aeschbacher and the rest of the Coop lab for helpful comments and discussion of the manuscript. This work was supported by grants from the National Science Foundation under Grant No. 1262645 to PLR and GC and by the National Institute of General Medical Sciences of the National Institutes of Health under award numbers NIH RO1GM83098 and RO1GM107374 to GC.

## Supporting Information

### The exact solution to the equilibrium frequency

Here we describe how to solve (24) exactly in one dimension, i.e., the Fisher-KPP equation with piecewise constant selection. Let *p*(*t, x*) be the proportion of the allele under spatially varying selection *s*(*x*) in one dimension, so that

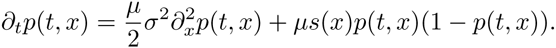

The stable distribution *Φ*(*x*) = lim_*t→∞*_ *p*(*t, x*) then solves

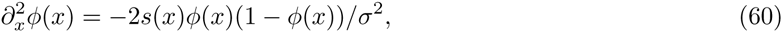

with appropriate boundary conditions. First rescale space by 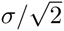 so the *σ*^2^*/*2 term disappears. If we now assume that *s*(*x*) is piecewise constant, *s*(*x*) = *s*_*i*_ for *x ∈* [*x*_*i*_*,x*_*i*+1_), with *x*_0_ = −*∞* and *x*_*n*+2_ = *∞*, then the equation is integrable: if we multiply through by 2*∂*_*x*_*ϕ*(*x*) and integrate, then we get that

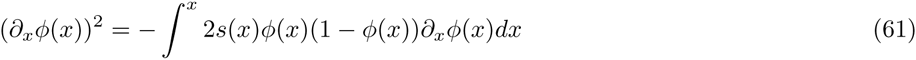

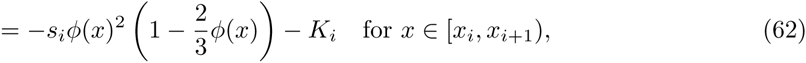

if we define

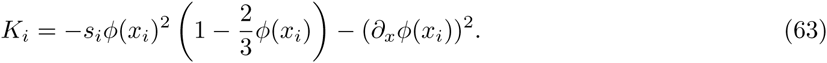

For ease of reference, we define

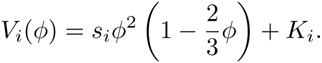

Note that 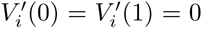, that *V*_*i*_(0) = *K*_*i*_ and *V*_*i*_(1) = *K*_*i*_ + *s*_*i*_*/*3. We will always have that *V* (*ϕ*) *≤* 0. (We have then that 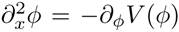, the equation of motion of a particle in potential *V*. Also note that this implies “conservation of energy”, i.e., (*∂*_*x*_*ϕ*(*x*))^2^ + *V* (*ϕ*(*x*)) is constant.) Rearranging, we get that 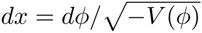, so where *ϕ*(*x*) is monotone, the inverse is

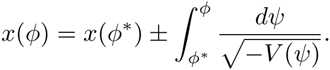

For each *i* then define the elliptic function

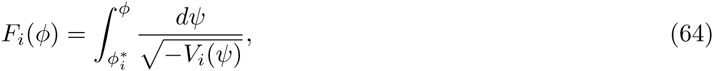

where take the positive branch of the square root, and 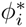 will be chosen later. Then we have that *x*(*ϕ*) − *x*(*ϕ*_0_) = *±*(*F* (*ϕ*) − *F* (*ϕ*_0_)), or for an appropriate *x*_0_,

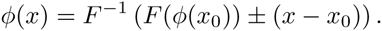

We clearly want lim_*x→ϕ*_ *∂*_*x*_*ϕ*(*±x*) = 0, and lim_*x→∞*_ *ϕ*(*±x*) to be zero or one depending on the sign of *s*_0_ and *s*_*n*+1_. Since (*∂*_*x*_*ϕ*(*x*))^2^ = −*V* (*x*), this implies that if *s*_0_ *<* 0, then *K*_0_ = 0, while if *s*_0_ *>* 0 then *K*_0_ = *s*_0_*/*6; and likewise for *K*_*n*+1_.

We also require that *ϕ*(*x*) and *ϕ*′ (*x*) are continuous. Continuity of *ϕ* (*x*) is equivalent to *V*_*i*_(*x*_*i*+1_) = *V*_*i*+1_(*x*_*i*+1_), which we can rearrange to find an equation for *K*_*i*+1_ in terms of *K*_*i*_ and *ϕ*(*x*_*i*+1_):

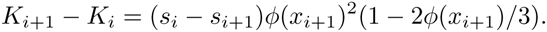

What about the frequency at the points the selection changes, *ϕ*(*x*_*i*_)? Well, if *ϕ*(*x*) is monotone on [*x*_*i*_*,x*_*i*+1_) then we can without loss of generality take 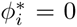 or 1 depending on the sign of *s*_*i*_. Otherwise, let 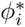 be the (unique) root of *V*_*i*_ in [*x*_*i*_*,x*_*i*+1_), so 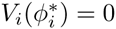. In this case, 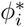 is the maximum or minimum of *ϕ* in the interval: if *s >* 0, then *ϕ*_*i*_ = max*{ϕ*(*x*) : *x ∈* [*x*_*i*_*,x*_*i*+1_)*}*. Recall we defined *F*_*i*_ using 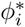; now using the fact that *ϕ* is monotone with the opposite sign on either side of 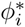, 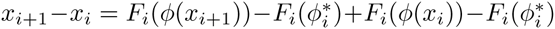, and that 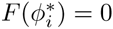, we know that the length of the *i*th stretch is

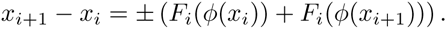

Note that all the *±*’s are easily relatable to the signs of *s*_*i*_. If we knew *ϕ*(*x* _1_) and *ϕ*′ (*x* _1_), then we’d be able to solve the equations for *ϕ*(*x*_*i*_) and *K*_*i*_ recursively upwards. In some cases, such as Slatkin [1973], we can infer *ϕ*(0) and *∂*_*x*_*ϕ*(0) by spatial symmetry. In other cases, we are only given *ϕ*(*−∞*) and *ϕ*(*∞*), and have to work inwards from the ends.

### Doing the integrals

An important ingredient in the above method is the integral (64), to which a method for solving the Korteweg-de Vries equation can be applied [NEQwiki, 2013]. Recall that *V*_*i*_(*ϕ*) = *s*_*i*_*ϕ*^2^(1 − 2*ϕ/*3) + *K*_*i*_, with *K*_*i*_ = *V*_*i*_(0) chosen to match *V* at the boundaries; since we’re just working within an interval with *s* constant, we can rescale space by 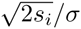, so that *s* and *σ* drop from the equation. We then want to integrate

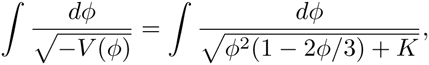

over a domain where *V* is always negative. Let *ϕ*^2^(1 − 2*ϕ/*3) + *K* = (*α − ϕ*)(*ϕ − β*)(*ϕ − γ*), and change variables first to *y*^2^ = (*α − ϕ*), and then to 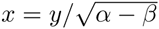, so that

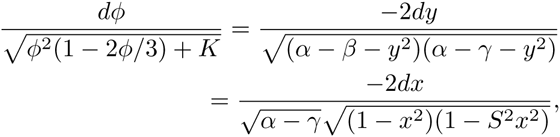

with *S*^2^ = (*α − β*)*/*(*α − γ*). Now Jacobi’s incomplete elliptic integral of the first kind is defined by

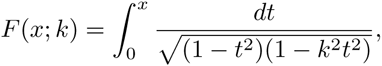

and the Jacobian elliptic function sn(*x*; *k*) is the inverse: *F* (sn(*x*; *k*); *k*) = *x*. As *k →* 0, sn(*x*; *k*) *→* sin(*x*), while as *k →* 1, sn(*x*; *k*) *→* sinh(*x*).

### More information about the simulations

Here we provide additional details regarding the simulations we used to validate the theory in figures 4 and 5.

**Figure 1.**
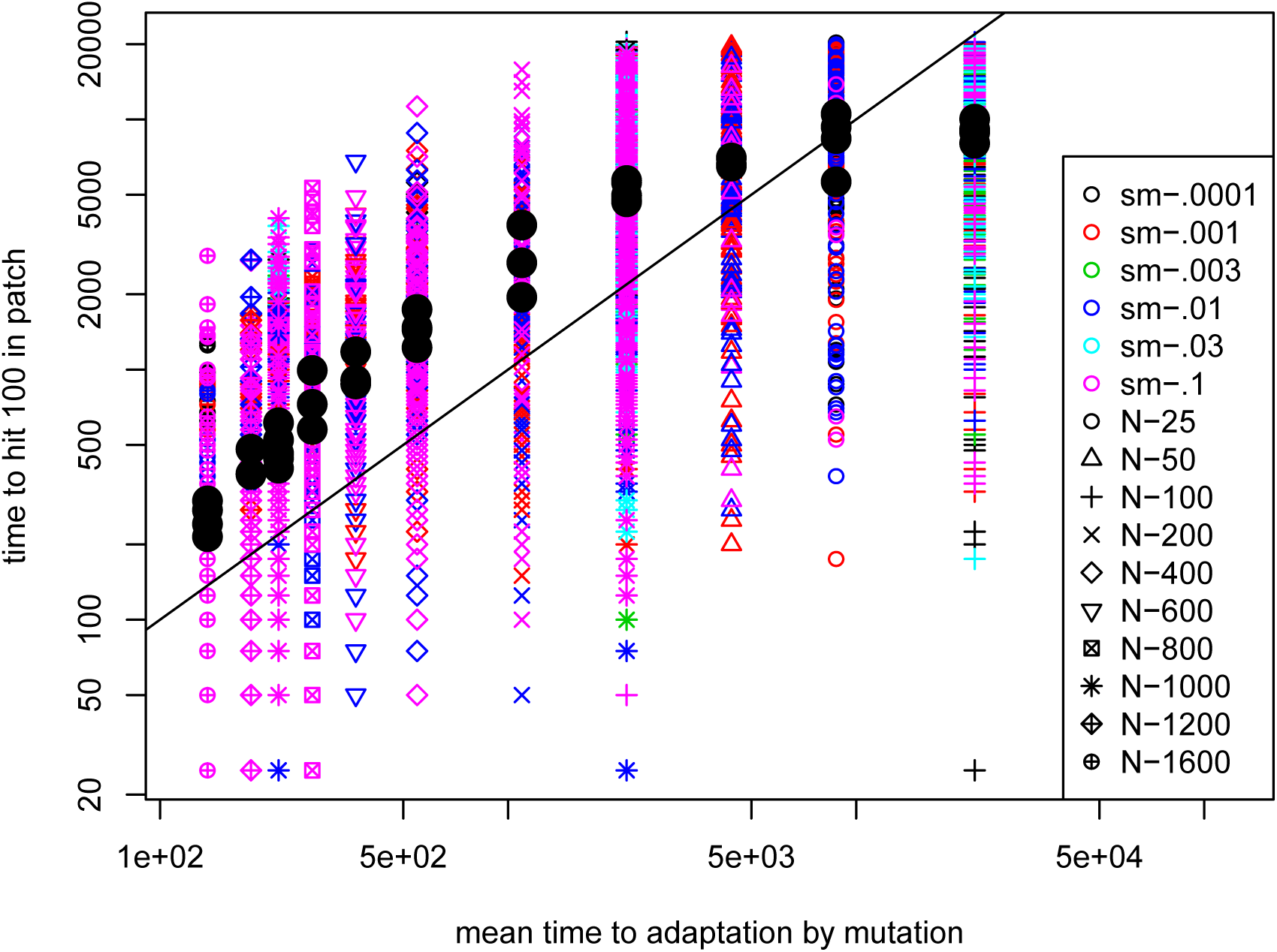
The same data shown in the left panel of figure 4, but all times shown (not just the interquartile ranges), and including those parameter values at which most of the simulations did not adapt by 25,000 generations.

**Figure 2.**
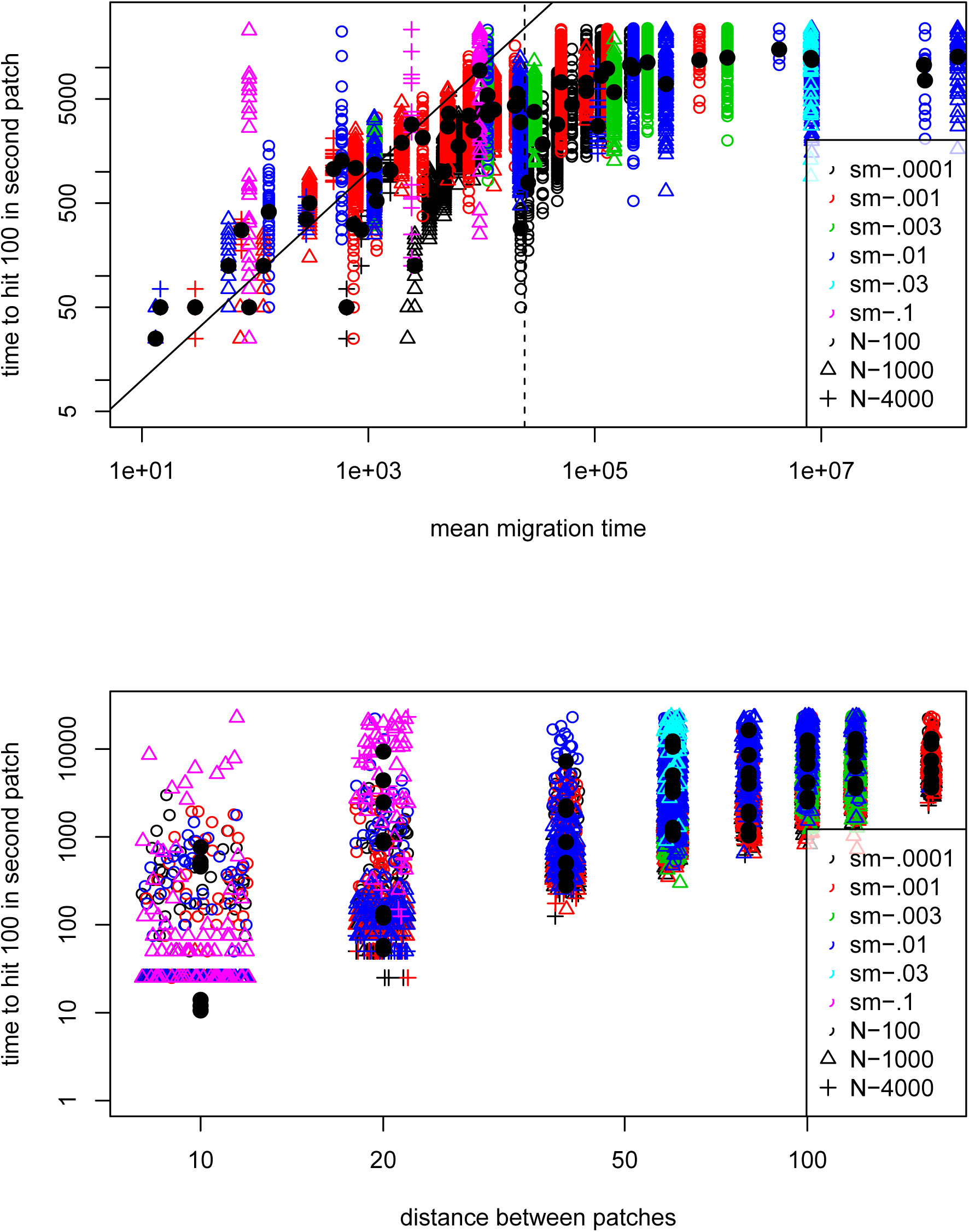
The same data shown in the right panel of figure 4, but all times shown (not just the interquartile ranges), and including those parameter values at which most of the simulations did not adapt by 25,000 generations. The upper panel has the predicted time to adaptation on the horizontal axis as in figure 4, and the lower panel has, for comparison, the raw distance between patches (which predicts time to adaptation, but not as well).

**Figure 3.**
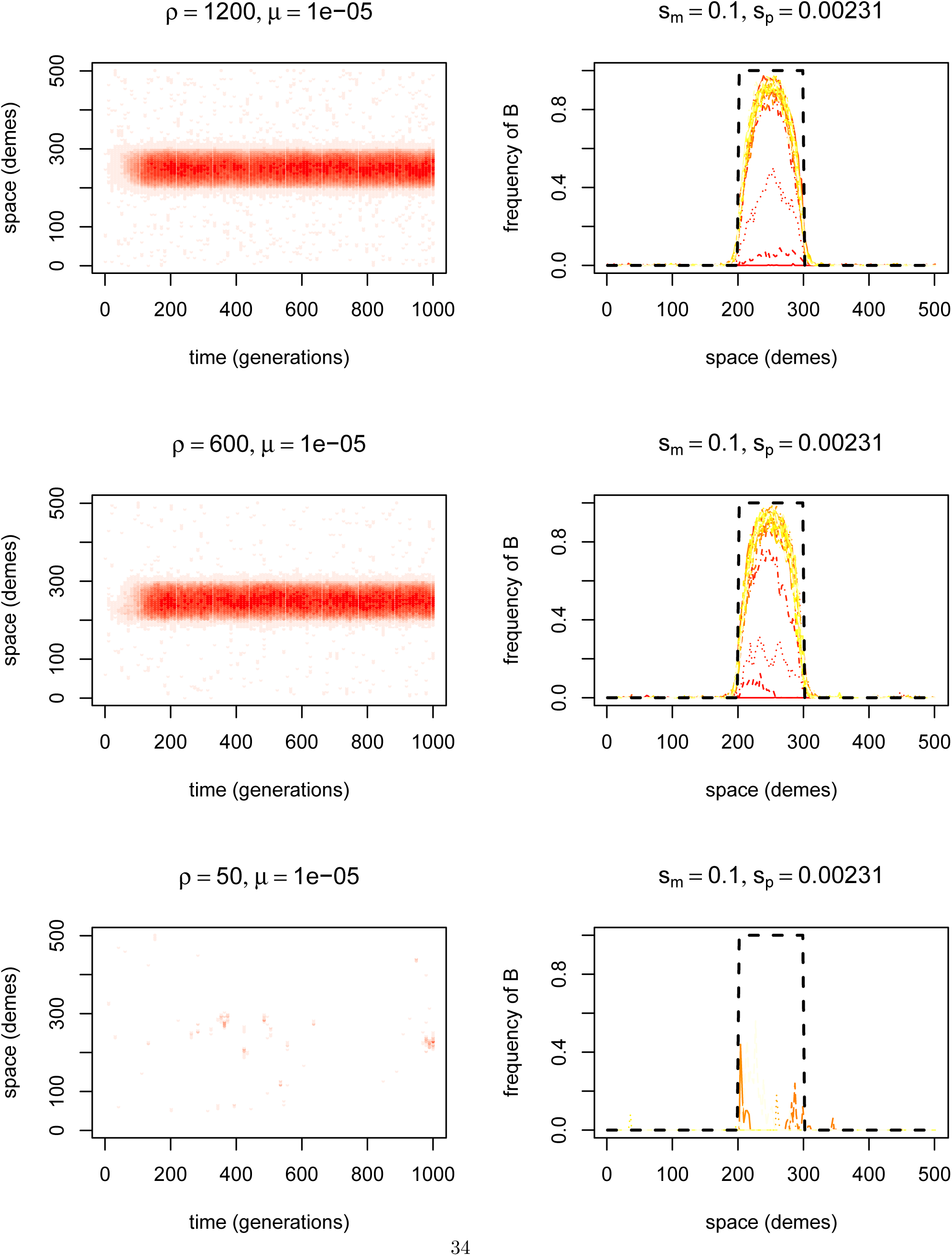
Randomly chosen simulations of adaptation by new mutation with *s*_*m*_ = 0.1, *σ ≈* 1, and *ρ* varying. On the left of each is a space-time heatmap of the local frequency of *B* alleles; and on the right are twenty-five curves showing the frequencies of *B* at evenly spaced time points (i.e., each line represents a vertical slice through the plot on the left); dotted black lines indicate the patches where *B* is advantageous.

**Figure 4.**
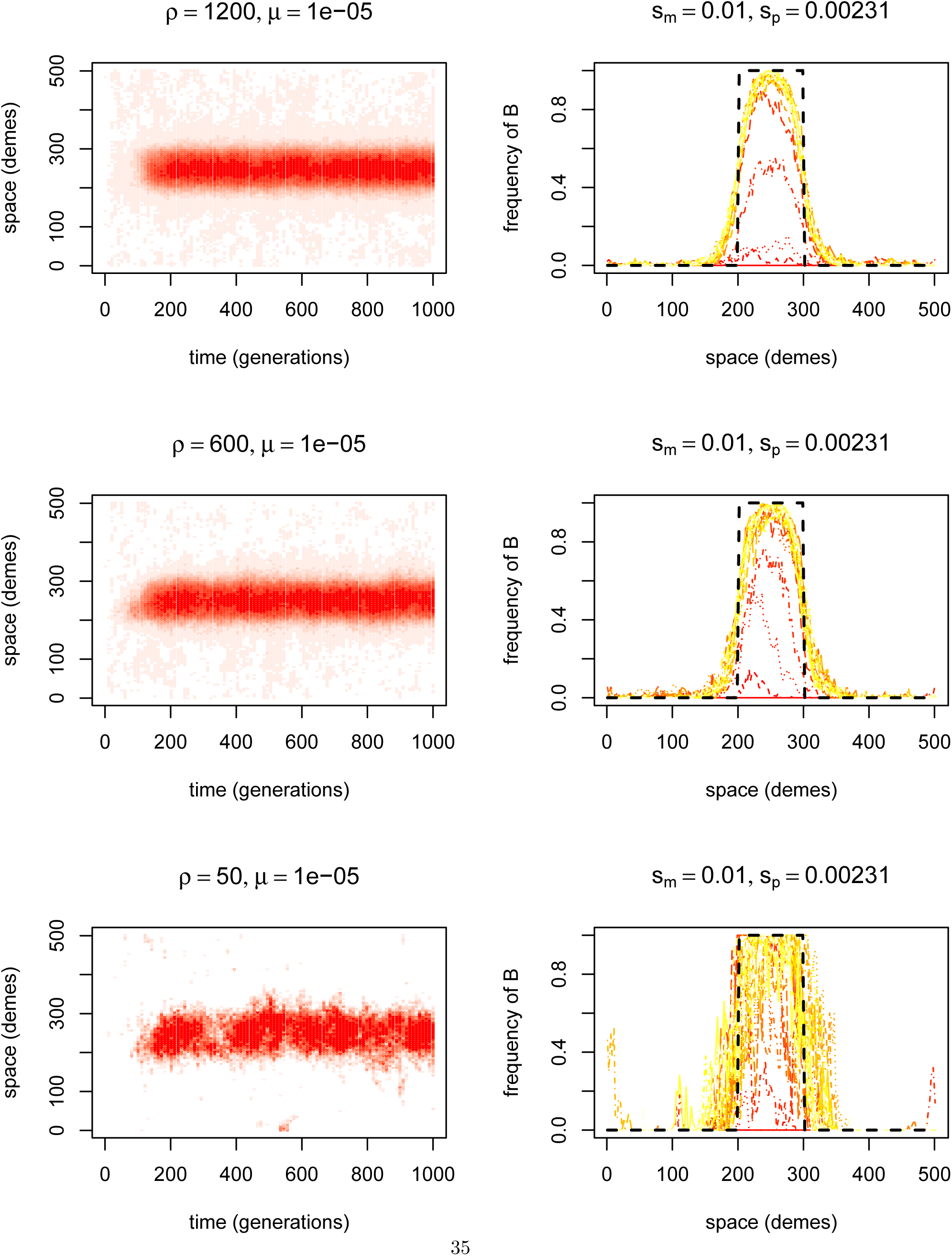
Randomly chosen simulations of adaptation by new mutation with *s*_*m*_ = 0.01, *σ ≈* 1, and *ρ* varying. On the left of each is a space-time heatmap of the local frequency of *B* alleles; and on the right are twenty-five curves showing the frequencies of *B* at evenly spaced time points (i.e., each line represents a vertical slice through the plot on the left); dotted black lines indicate the patches where *B* is advantageous.

**Figure 5.**
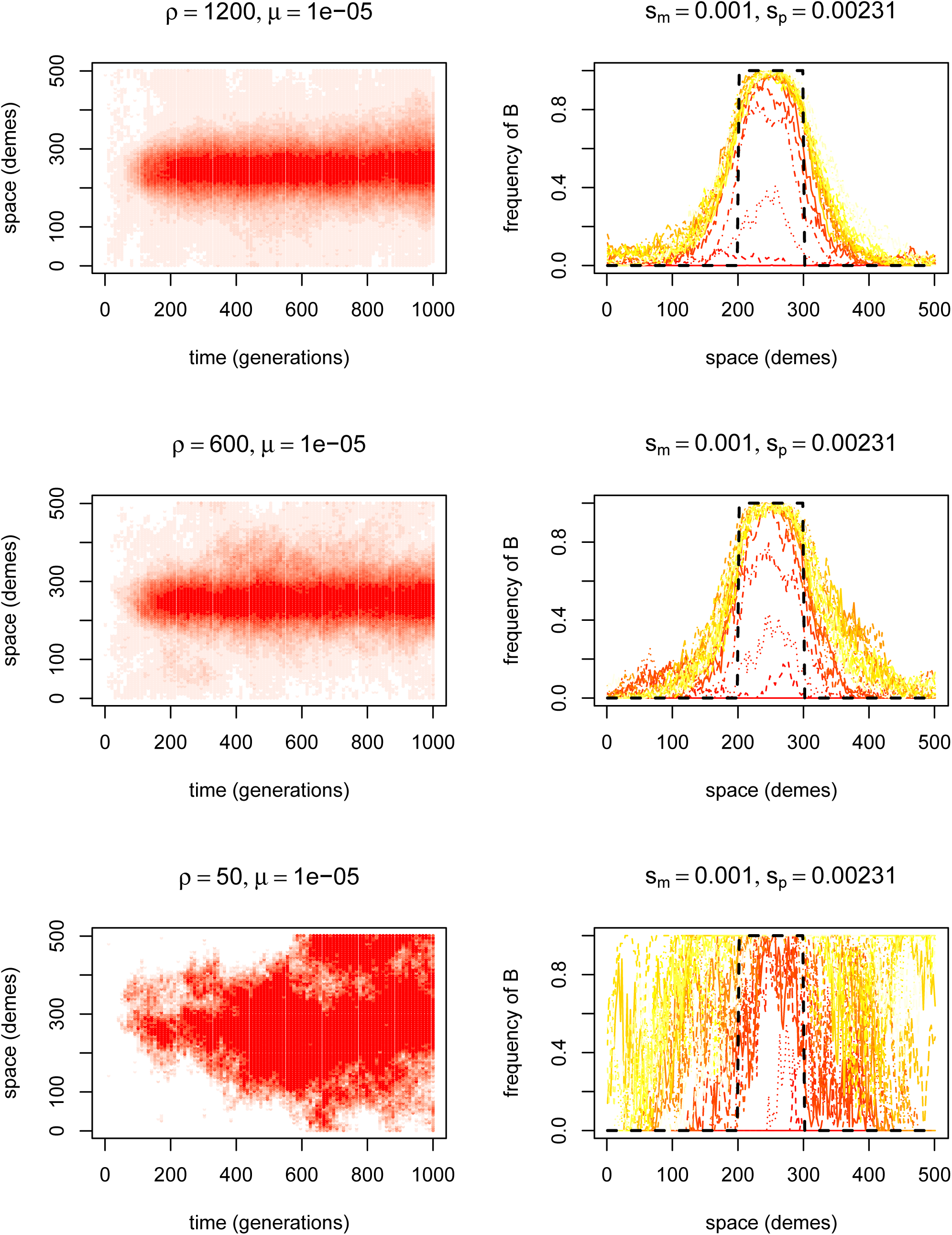
Randomly chosen simulations of adaptation by new mutation with *s*_*m*_ = 0.001, *σ ≈* 1, and *ρ* varying. On the left of each is a space-time heatmap of the local frequency of *B* alleles; and on the right are twenty-five curves showing the frequencies of *B* at evenly spaced time points (i.e., each line represents a vertical slice through the plot on the left); dotted black lines indicate the patches where *B* is advantageous.

**Figure 6.**
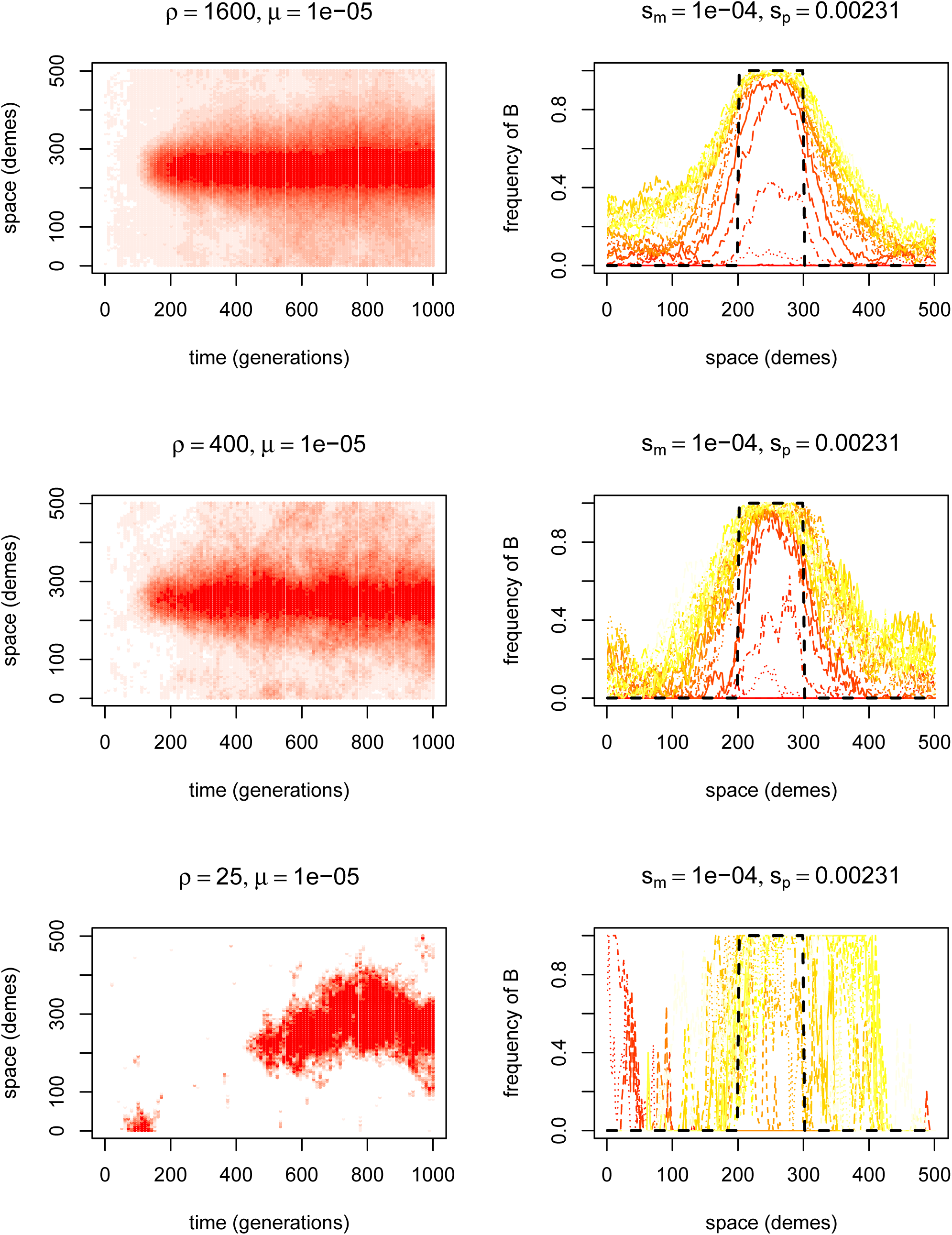
Randomly chosen simulations of adaptation by new mutation with *s*_*m*_ = 0.001, *σ ≈* 1, and *ρ* varying. On the left of each is a space-time heatmap of the local frequency of *B* alleles; and on the right are twenty-five curves showing the frequencies of *B* at evenly spaced time points (i.e., each line represents a vertical slice through the plot on the left); dotted black lines indicate the patches where *B* is advantageous.

**Figure 7.**
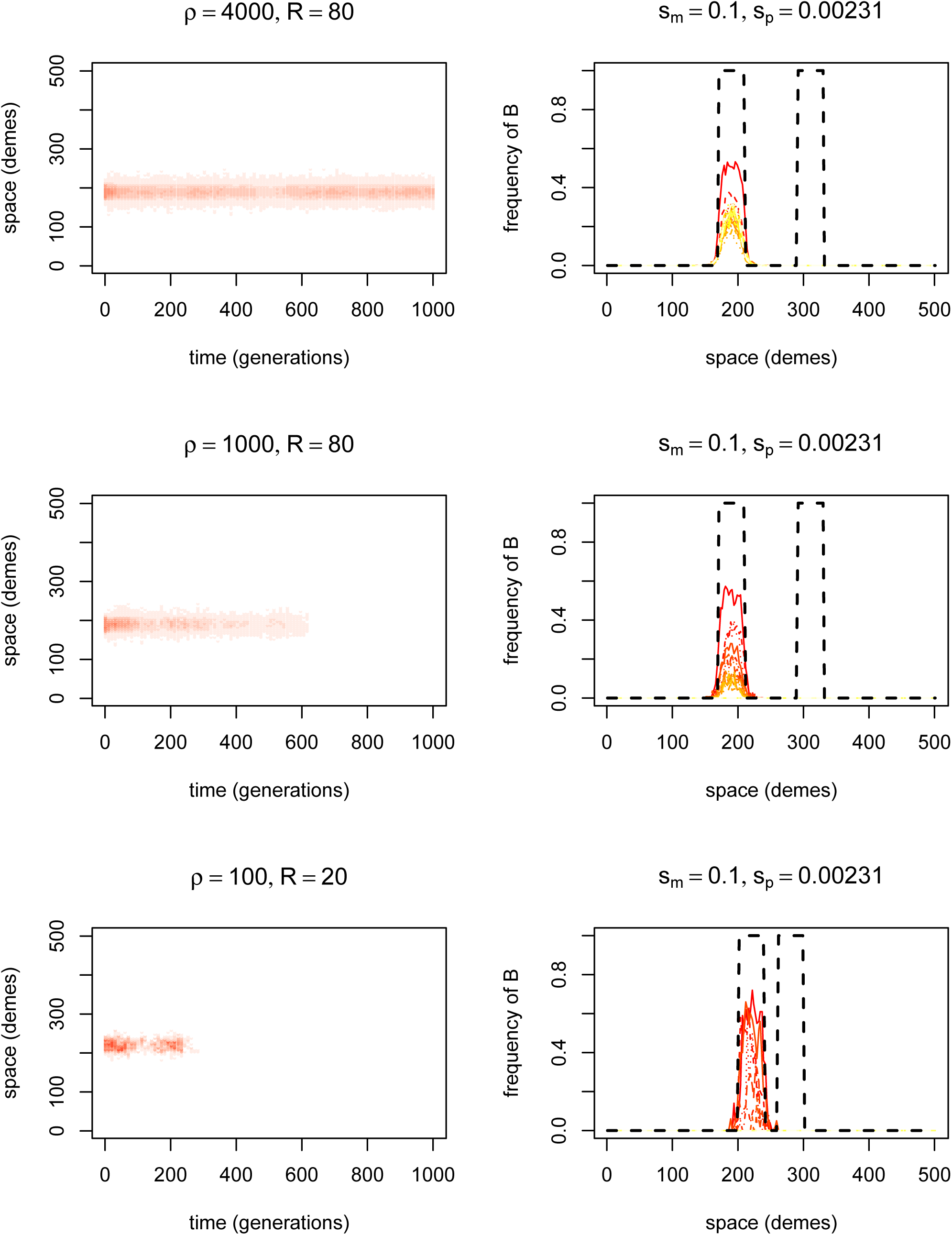
Randomly chosen simulations of adaptation by migration with *s*_*m*_ = 0.1, *R* = 80, *σ ≈* 1, and *ρ* varying. On the left of each is a space-time heatmap of the local frequency of *B* alleles; and on the right are twenty-five curves showing the frequencies of *B* at evenly spaced time points (i.e., each line represents a vertical slice through the plot on the left); dotted black lines indicate the patches where *B* is advantageous.

**Figure 8.**
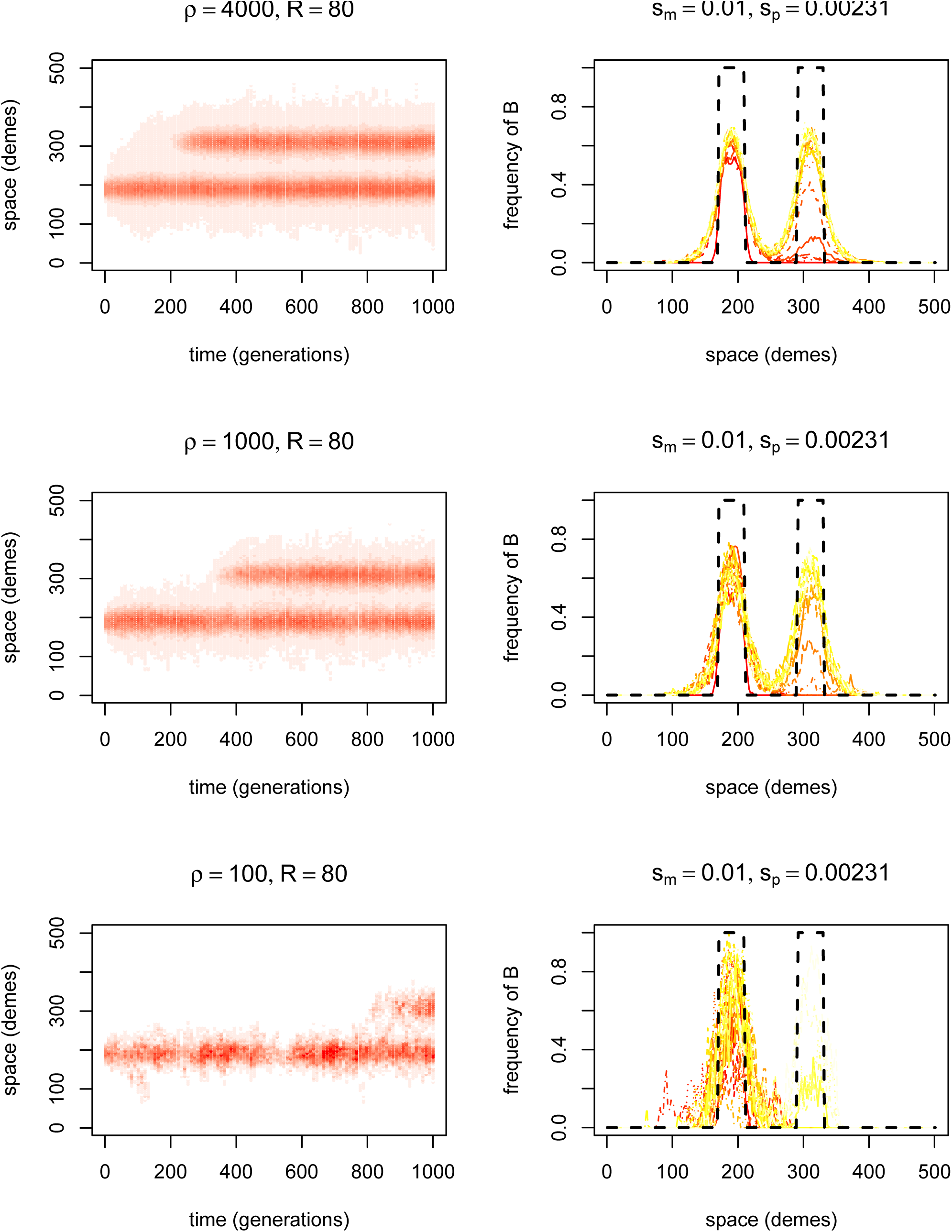
Randomly chosen simulations of adaptation by migration with *s*_*m*_ = 0.01, *R* = 80, *σ ≈* 1, and *ρ* varying. On the left of each is a space-time heatmap of the local frequency of *B* alleles; and on the right are twenty-five curves showing the frequencies of *B* at evenly spaced time points (i.e., each line represents a vertical slice through the plot on the left); dotted black lines indicate the patches where *B* is advantageous.

**Figure 9.**
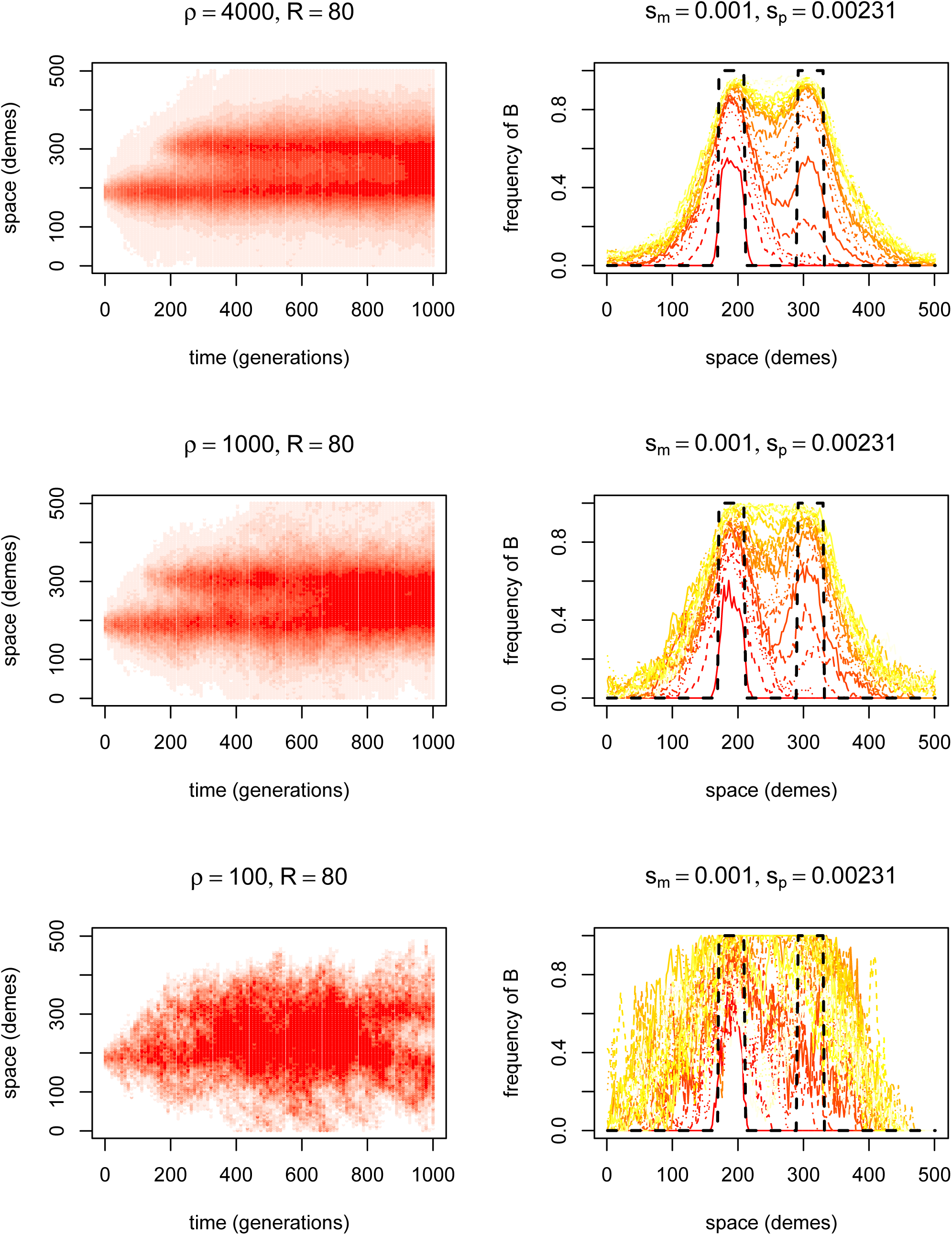
Randomly chosen simulations of adaptation by migration with *s*_*m*_ = 0.001, *R* = 80, *σ ≈* 1, and *ρ* varying. On the left of each is a space-time heatmap of the local frequency of *B* alleles; and on the right are twenty-five curves showing the frequencies of *B* at evenly spaced time points (i.e., each line represents a vertical slice through the plot on the left); dotted black lines indicate the patches where *B* is advantageous.

**Figure 10.**
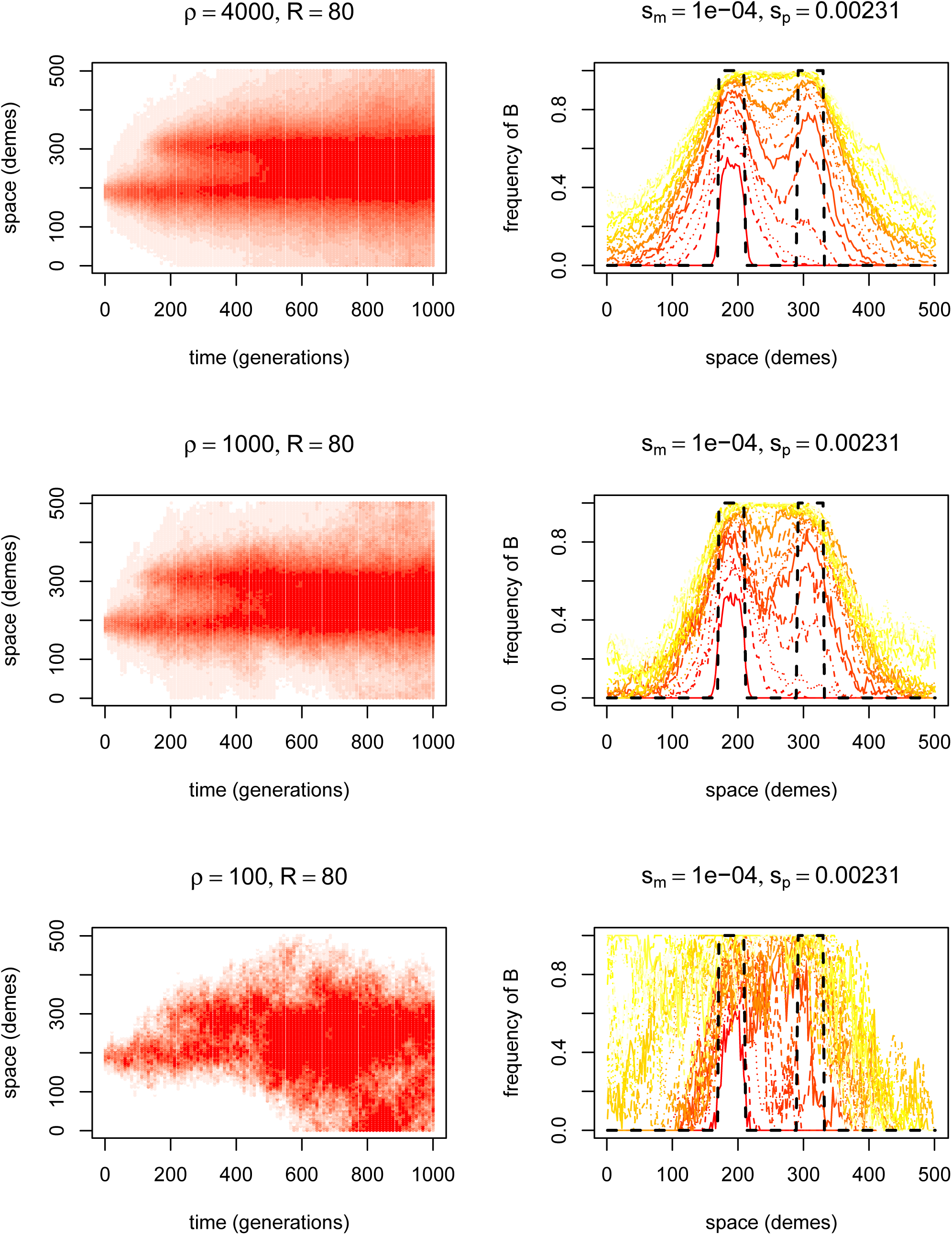
Randomly chosen simulations of adaptation by migration with *s*_*m*_ = 0.0001, *R* = 80, *σ ≈* 1, and *ρ* varying. On the left of each is a space-time heatmap of the local frequency of *B* alleles; and on the right are twenty-five curves showing the frequencies of *B* at evenly spaced time points (i.e., each line represents a vertical slice through the plot on the left); dotted black lines indicate the patches where *B* is

**Table 1.** Parameter values used in estimates of mean time to adaptation by mutation of figure 4. All simulations also used a linear grid of 501 demes with a patch of 99 demes in the center, the migration model described in Simulation methods, *μ* = 10^−5^, and *s*_*p*_ = .0023 (calculated as the growth rate as described in the text). *T*adapt is the mean time until 100 *B* alleles were present in the patch, and *p*adapted is the proportion of the simulations that adapted by the 25,000 generations. The simulation began with no *B* alleles.

| $\mu$ | $\rho$ | $s_p$ | $s_m$ | $T_{\text{adapt}}$ | $p_{\text{adapted}}$ | n | $\mu$ | $\rho$ | $s_p$ | $s_m$ | $T_{\text{adapt}}$ | $p_{\text{adapted}}$ | n |
| --- | --- | --- | --- | --- | --- | --- | --- | --- | --- | --- | --- | --- | --- |
| 1e-05 | 25 | 0.01 | -0.10 | 19630 | 0.06 | 100 | 1e-05 | 600 | 0.01 | -0.10 | 1082 | 1.00 | 40 |
| 1e-05 | 25 | 0.01 | -0.01 | 15533 | 0.55 | 100 | 1e-05 | 600 | 0.01 | -0.01 | 924 | 1.00 | 40 |
| 1e-05 | 25 | 0.01 | -0.00 | 14353 | 0.72 | 100 | 1e-05 | 600 | 0.01 | -0.01 | 828 | 1.00 | 40 |
| 1e-05 | 25 | 0.01 | -0.00 | 13282 | 0.74 | 100 | 1e-05 | 600 | 0.01 | -0.00 | 846 | 1.00 | 40 |
| 1e-05 | 50 | 0.01 | -0.10 | 11587 | 0.33 | 40 | 1e-05 | 600 | 0.01 | -0.00 | 966 | 1.00 | 40 |
| 1e-05 | 50 | 0.01 | -0.10 | 11586 | 0.38 | 40 | 1e-05 | 800 | 0.01 | -0.10 | 802 | 1.00 | 40 |
| 1e-05 | 50 | 0.01 | -0.01 | 9419 | 0.88 | 40 | 1e-05 | 800 | 0.01 | -0.10 | 1182 | 1.00 | 40 |
| 1e-05 | 50 | 0.01 | -0.01 | 8848 | 0.82 | 40 | 1e-05 | 800 | 0.01 | -0.01 | 846 | 1.00 | 40 |
| 1e-05 | 50 | 0.01 | -0.00 | 6529 | 0.95 | 40 | 1e-05 | 800 | 0.01 | -0.01 | 607 | 1.00 | 40 |
| 1e-05 | 50 | 0.01 | -0.00 | 9986 | 0.90 | 40 | 1e-05 | 800 | 0.01 | -0.00 | 646 | 1.00 | 40 |
| 1e-05 | 100 | 0.01 | -0.10 | 7425 | 0.91 | 316 | 1e-05 | 800 | 0.01 | -0.00 | 507 | 1.00 | 40 |
| 1e-06 | 100 | 0.01 | -0.10 | 21010 | 0.23 | 217 | 1e-05 | 1000 | 0.01 | -0.10 | 615 | 1.00 | 256 |
| 1e-05 | 100 | 0.01 | -0.03 | 6036 | 0.98 | 195 | 2.4e-06 | 1000 | 0.01 | -0.10 | 5309 | 0.96 | 257 |
| 1e-06 | 100 | 0.01 | -0.03 | 19859 | 0.31 | 194 | 1e-05 | 1000 | 0.01 | -0.03 | 523 | 1.00 | 194 |
| 1e-05 | 100 | 0.01 | -0.01 | 5408 | 0.98 | 315 | 1e-06 | 1000 | 0.01 | -0.03 | 6425 | 0.95 | 195 |
| 1e-06 | 100 | 0.01 | -0.01 | 20606 | 0.28 | 216 | 1e-05 | 1000 | 0.01 | -0.01 | 449 | 1.00 | 254 |
| 1e-05 | 100 | 0.01 | -0.00 | 4745 | 0.99 | 195 | 2.4e-06 | 1000 | 0.01 | -0.01 | 4806 | 0.96 | 256 |
| 1e-06 | 100 | 0.01 | -0.00 | 19564 | 0.33 | 195 | 1e-05 | 1000 | 0.01 | -0.00 | 460 | 1.00 | 193 |
| 1e-05 | 100 | 0.01 | -0.00 | 4894 | 0.99 | 315 | 1e-06 | 1000 | 0.01 | -0.00 | 5203 | 0.98 | 194 |
| 1e-06 | 100 | 0.01 | -0.00 | 19417 | 0.33 | 216 | 1e-05 | 1000 | 0.01 | -0.00 | 444 | 1.00 | 255 |
| 1e-05 | 100 | 0.01 | -0.00 | 4940 | 0.99 | 316 | 2.4e-06 | 1000 | 0.01 | -0.00 | 4037 | 1.00 | 257 |
| 1e-06 | 100 | 0.01 | -0.00 | 18025 | 0.40 | 217 | 1e-05 | 1000 | 0.01 | -0.00 | 404 | 1.00 | 216 |
| 1e-05 | 200 | 0.01 | -0.10 | 4296 | 0.97 | 40 | 1e-06 | 1000 | 0.01 | -0.00 | 4567 | 1.00 | 216 |
| 1e-05 | 200 | 0.01 | -0.10 | 3736 | 1.00 | 40 | 1e-05 | 1200 | 0.01 | -0.10 | 501 | 1.00 | 40 |
| 1e-05 | 200 | 0.01 | -0.01 | 2943 | 1.00 | 40 | 1e-05 | 1200 | 0.01 | -0.10 | 460 | 1.00 | 40 |
| 1e-05 | 200 | 0.01 | -0.01 | 2402 | 1.00 | 40 | 1e-05 | 1200 | 0.01 | -0.01 | 336 | 1.00 | 40 |
| 1e-05 | 200 | 0.01 | -0.00 | 1899 | 1.00 | 40 | 1e-05 | 1200 | 0.01 | -0.01 | 431 | 1.00 | 40 |
| 1e-05 | 200 | 0.01 | -0.00 | 1992 | 1.00 | 40 | 1e-05 | 1200 | 0.01 | -0.00 | 390 | 1.00 | 40 |
| 1e-05 | 400 | 0.01 | -0.10 | 1740 | 1.00 | 100 | 1e-05 | 1200 | 0.01 | -0.00 | 372 | 1.00 | 40 |
| 1e-05 | 400 | 0.01 | -0.01 | 1471 | 1.00 | 100 | 1e-05 | 1600 | 0.01 | -0.10 | 299 | 1.00 | 100 |
| 1e-05 | 400 | 0.01 | -0.00 | 1446 | 1.00 | 100 | 1e-05 | 1600 | 0.01 | -0.01 | 462 | 0.99 | 100 |
| 1e-05 | 400 | 0.01 | -0.00 | 1224 | 1.00 | 100 | 1e-05 | 1600 | 0.01 | -0.00 | 241 | 1.00 | 100 |
| 1e-05 | 600 | 0.01 | -0.10 | 1276 | 1.00 | 40 | 1e-05 | 1600 | 0.01 | -0.00 | 274 | 1.00 | 100 |

**Table 2.** Parameter values used in estimates of mean time to adaptation by migration of figure 4. All simulations also used a linear grid of 501 demes with two patches of 99 demes each, separated by *R* demes in the center; the migration model described in Simulation methods, *μ* = 10^−5^, and *s*_*p*_ = .0023 (calculated as the growth rate as described in the text). *T*adapt is the mean time until 100 *B* alleles were present in the patch, and *p*adapted is the proportion of the simulations that adapted by the 25,000 generations. At the start of the simulation, one patch was initialized with a frequency of 0.8 *B* alleles, which were absent elsewhere.

| R | $\rho$ | $s_p$ | $s_m$ | $T_{\text{adapt}}$ | $p_{\text{adapted}}$ | n | R | $\rho$ | $s_p$ | $s_m$ | $T_{\text{adapt}}$ | $p_{\text{adapted}}$ | n |
| --- | --- | --- | --- | --- | --- | --- | --- | --- | --- | --- | --- | --- | --- |
| 100 | 100 | 0.01 | -1e-01 | 24975 | 0.00 | 300 | 60 | 1000 | 0.01 | -3e-02 | 22224 | 0.21 | 200 |
| 10 | 100 | 0.01 | -1e-01 | 24975 | 0.00 | 40 | 100 | 1000 | 0.01 | -1e-02 | 18098 | 0.55 | 299 |
| 120 | 100 | 0.01 | -1e-01 | 24975 | 0.00 | 295 | 10 | 1000 | 0.01 | -1e-02 | 14 | 1.00 | 100 |
| 160 | 100 | 0.01 | -1e-01 | 24975 | 0.00 | 40 | 120 | 1000 | 0.01 | -1e-02 | 23242 | 0.14 | 297 |
| 20 | 100 | 0.01 | -1e-01 | 24975 | 0.00 | 40 | 160 | 1000 | 0.01 | -1e-02 | 24762 | 0.01 | 100 |
| 40 | 100 | 0.01 | -1e-01 | 24975 | 0.00 | 40 | 20 | 1000 | 0.01 | -1e-02 | 136 | 1.00 | 100 |
| 60 | 100 | 0.01 | -1e-01 | 24975 | 0.00 | 300 | 40 | 1000 | 0.01 | -1e-02 | 874 | 1.00 | 99 |
| 80 | 100 | 0.01 | -1e-01 | 24975 | 0.00 | 40 | 60 | 1000 | 0.01 | -1e-02 | 3485 | 1.00 | 300 |
| 100 | 100 | 0.01 | -3e-02 | 24975 | 0.00 | 200 | 80 | 1000 | 0.01 | -1e-02 | 9204 | 0.96 | 100 |
| 120 | 100 | 0.01 | -3e-02 | 24975 | 0.00 | 199 | 100 | 1000 | 0.01 | -3e-03 | 4168 | 1.00 | 200 |
| 60 | 100 | 0.01 | -3e-02 | 24903 | 0.01 | 200 | 120 | 1000 | 0.01 | -3e-03 | 6304 | 1.00 | 200 |
| 100 | 100 | 0.01 | -1e-02 | 24354 | 0.04 | 300 | 60 | 1000 | 0.01 | -3e-03 | 1273 | 1.00 | 199 |
| 10 | 100 | 0.01 | -1e-02 | 1705 | 0.95 | 40 | 100 | 1000 | 0.01 | -1e-03 | 2813 | 1.00 | 299 |
| 120 | 100 | 0.01 | -1e-02 | 24802 | 0.01 | 299 | 10 | 1000 | 0.01 | -1e-03 | 12 | 1.00 | 100 |
| 160 | 100 | 0.01 | -1e-02 | 24975 | 0.00 | 40 | 120 | 1000 | 0.01 | -1e-03 | 4002 | 1.00 | 300 |
| 20 | 100 | 0.01 | -1e-02 | 3601 | 0.95 | 40 | 160 | 1000 | 0.01 | -1e-03 | 7386 | 1.00 | 100 |
| 40 | 100 | 0.01 | -1e-02 | 9463 | 0.88 | 40 | 20 | 1000 | 0.01 | -1e-03 | 121 | 1.00 | 100 |
| 60 | 100 | 0.01 | -1e-02 | 18674 | 0.45 | 299 | 40 | 1000 | 0.01 | -1e-03 | 507 | 1.00 | 100 |
| 80 | 100 | 0.01 | -1e-02 | 23899 | 0.12 | 40 | 60 | 1000 | 0.01 | -1e-03 | 1115 | 1.00 | 300 |
| 100 | 100 | 0.01 | -3e-03 | 14601 | 0.80 | 200 | 80 | 1000 | 0.01 | -1e-03 | 1941 | 1.00 | 100 |
| 120 | 100 | 0.01 | -3e-03 | 18645 | 0.53 | 199 | 100 | 1000 | 0.01 | -1e-04 | 2528 | 1.00 | 300 |
| 60 | 100 | 0.01 | -3e-03 | 5424 | 0.98 | 200 | 10 | 1000 | 0.01 | -1e-04 | 11 | 1.00 | 99 |
| 100 | 100 | 0.01 | -1e-03 | 8992 | 0.96 | 299 | 120 | 1000 | 0.01 | -1e-04 | 3590 | 1.00 | 298 |
| 10 | 100 | 0.01 | -1e-03 | 522 | 1.00 | 40 | 160 | 1000 | 0.01 | -1e-04 | 5648 | 1.00 | 100 |
| 120 | 100 | 0.01 | -1e-03 | 11375 | 0.93 | 300 | 20 | 1000 | 0.01 | -1e-04 | 122 | 1.00 | 100 |
| 160 | 100 | 0.01 | -1e-03 | 16930 | 0.68 | 40 | 40 | 1000 | 0.01 | -1e-04 | 498 | 1.00 | 100 |
| 20 | 100 | 0.01 | -1e-03 | 1518 | 0.97 | 40 | 60 | 1000 | 0.01 | -1e-04 | 1051 | 1.00 | 298 |
| 40 | 100 | 0.01 | -1e-03 | 2234 | 1.00 | 40 | 80 | 1000 | 0.01 | -1e-04 | 1796 | 1.00 | 100 |
| 60 | 100 | 0.01 | -1e-03 | 4277 | 0.98 | 300 | 10 | 4000 | 0.01 | -1e-01 | 0 | 1.00 | 20 |
| 80 | 100 | 0.01 | -1e-03 | 5328 | 1.00 | 40 | 160 | 4000 | 0.01 | -1e-01 | 24975 | 0.00 | 20 |
| 100 | 100 | 0.01 | -1e-04 | 6867 | 0.99 | 299 | 20 | 4000 | 0.01 | -1e-01 | 4406 | 1.00 | 20 |
| 10 | 100 | 0.01 | -1e-04 | 461 | 1.00 | 40 | 40 | 4000 | 0.01 | -1e-01 | 24975 | 0.00 | 20 |
| 120 | 100 | 0.01 | -1e-04 | 9347 | 0.99 | 300 | 80 | 4000 | 0.01 | -1e-01 | 24975 | 0.00 | 20 |
| 160 | 100 | 0.01 | -1e-04 | 12731 | 0.90 | 40 | 10 | 4000 | 0.01 | -1e-02 | 3746 | 0.85 | 20 |
| 20 | 100 | 0.01 | -1e-04 | 860 | 1.00 | 40 | 160 | 4000 | 0.01 | -1e-02 | 24975 | 0.00 | 20 |
| 40 | 100 | 0.01 | -1e-04 | 2031 | 1.00 | 40 | 20 | 4000 | 0.01 | -1e-02 | 58 | 1.00 | 20 |
| 60 | 100 | 0.01 | -1e-04 | 3529 | 0.99 | 300 | 40 | 4000 | 0.01 | -1e-02 | 362 | 1.00 | 20 |
| 80 | 100 | 0.01 | -1e-04 | 4839 | 1.00 | 40 | 80 | 4000 | 0.01 | -1e-02 | 4025 | 1.00 | 20 |
| 100 | 1000 | 0.01 | -1e-01 | 24975 | 0.00 | 300 | 10 | 4000 | 0.01 | -1e-03 | 6244 | 0.75 | 20 |
| 10 | 1000 | 0.01 | -1e-01 | 1004 | 0.99 | 100 | 160 | 4000 | 0.01 | -1e-03 | 4399 | 1.00 | 20 |
| 120 | 1000 | 0.01 | -1e-01 | 24975 | 0.00 | 298 | 20 | 4000 | 0.01 | -1e-03 | 52 | 1.00 | 20 |
| 160 | 1000 | 0.01 | -1e-01 | 24975 | 0.00 | 100 | 40 | 4000 | 0.01 | -1e-03 | 281 | 1.00 | 20 |
| 20 | 1000 | 0.01 | -1e-01 | 18403 | 0.42 | 100 | 80 | 4000 | 0.01 | -1e-03 | 1209 | 1.00 | 20 |
| 40 | 1000 | 0.01 | -1e-01 | 24965 | 0.00 | 100 | 10 | 4000 | 0.01 | -1e-04 | 3746 | 0.85 | 20 |
| 60 | 1000 | 0.01 | -1e-01 | 24975 | 0.00 | 300 | 160 | 4000 | 0.01 | -1e-04 | 3798 | 1.00 | 20 |
| 80 | 1000 | 0.01 | -1e-01 | 24975 | 0.00 | 100 | 20 | 4000 | 0.01 | -1e-04 | 52 | 1.00 | 20 |
| 100 | 1000 | 0.01 | -3e-02 | 24975 | 0.00 | 200 | 40 | 4000 | 0.01 | -1e-04 | 279 | 1.00 | 20 |
| 120 | 1000 | 0.01 | -3e-02 | 24975 | 0.00 | 200 | 80 | 4000 | 0.01 | -1e-04 | 1039 | 1.00 | 20 |

